# Archaeal SepF is essential for cell division in *Haloferax volcanii*

**DOI:** 10.1101/2020.10.06.327809

**Authors:** Phillip Nußbaum, Maren Gerstner, Marie Dingethal, Celine Erb, Sonja-Verena Albers

## Abstract

Bacterial cell division has been studied for decades but reports on the different archaeal cell division systems are rare. In many archaea, cell division depends on the tubulin homolog FtsZ, but further components of the divisome in these archaea are unknown. The halophilic archaeon *Haloferax volcanii* encodes two FtsZ homologs with different functions in cell division and a putative SepF homolog. In bacteria, SepF is part of the divisome and is recruited early to the FtsZ ring, where it most likely stimulates FtsZ ring formation. *H. volcanii* SepF co-localized with FtsZ1 and FtsZ2 at midcell. Overexpression of SepF had no effect on cell morphology, but no *sepF* deletion mutants could be generated. SepF depletion led to a severe cell division defect, resulting in cells with a strongly increased size. Overexpression of FtsZ1- and FtsZ2-GFP in SepF-depleted cells resulted in filamentous cells with an increasing number of FtsZ1 rings depending on the cell length, whereas FtsZ2 rings were not increased. Pull-down assays with HA-tagged SepF identified an interaction with FtsZ2 but not with FtsZ1. Archaeal SepF homologs lack the conserved glycine residue important for polymerization in bacteria and the *H. volcanii* SepF was purified as a dimer, suggesting that in contrast to the bacterial SepF homologs, polymerization does not seem to be important for its function. A model is proposed where first the FtsZ1 ring is formed and where SepF recruits FtsZ2 to the FtsZ1 ring, resulting in the formation of the FtsZ2 ring. This study provides important novel insights into cell division in archaea and shows that SepF is an important part of the divisome in FtsZ containing archaea.

## Introduction

Most archaeal and bacterial cells divide by binary fission, resulting in two daughter cells of equal size and DNA content. Cell division in most bacteria, and some archaea groups, starts with the polymerization of the conserved tubulin homologue FtsZ into a ring-like structure at midcell. Polymerization of FtsZ proteins into single stranded protofilaments is initiated after GTP binding (1). The so called Z-ring is associated with the cell membrane, providing a scaffold for down-stream cell division proteins, many of them involved in septal peptidoglycan synthesis (2–5). However, FtsZ lacks a membrane targeting domain and is therefore dependent on proteins that tether the FtsZ polymers to the membrane. In *Escherichia coli* FtsA and ZipA are responsible for the membrane association of FtsZ filaments. ZipA contains an N-terminal trans-membrane domain and is unique to Gammaproteobacteria (6). Like ZipA, FtsA binds to the conserved C-terminal peptide of FtsZ (7–9). FtsA, a protein that forms actin-like protofilaments in the presence of ATP, binds to the membrane via a C-terminal amphipathic helix (9–11). *Bacillus subtilis* lacks ZipA but has another FtsZ membrane anchor protein called SepF. SepF is highly conserved in Firmicutes, Actinobacteria and Cyanobacteria and acts as an alternative membrane anchor for FtsZ with an N-terminal amphipathic helix (12, 13). Yet, SepF binds to the conserved C-terminal peptide of FtsZ promoting the assembly and bundling of FtsZ polymers (14–16). A knock-out of *sepF* in *B. subtilis* resulted in deformed division septa which led to the proposal that SepF is required for a late step in cell division (13). This suggests that, besides anchoring FtsZ filaments to the membrane, SepF also has a regulatory role in cell division. Indeed it was discovered that the overexpression of SepF in *B. subtilis* leads to complete delocalization of late cell division proteins (17) and SepF overproduction in *Mycobacterium smegmatis* led to filamentation of the cells (18).

Electron microscopy of purified SepF from *B. subtilis* showed polymerization into large ring-like structures with an average diameter of about 50 nm *in vitro* (19). The observation of liposomes bound exclusively to the inside of SepF rings led to the assumption that *in vivo* the protein does not form rings but arcs, perpendicularly associated to the nascent septum (12). The ability of SepF to form rings *in vitro* is conserved in bacteria. A recent investigation of SepF homologs from different bacteria indicated that the SepF ring diameter correlates with the septum width (20), supporting the model that SepF forms a clamp on top of the leading edge of the growing septum. The crystal structure of SepF showed a dimer of two SepF monomers that contain a compact α/β-sandwich of two α-helices stacked against a five-stranded β-sheet. Dimers are formed by interaction of the β-sheets and polymerization is acquired by interaction of the α-helices of adjacent dimers (12). The highly conserved residue G109 of *B*.*subtillis* is important for interaction between dimers since a mutation in this residue leads to SepF dimers that do not polymerize (12). However, a recent study of SepF from *Corynebacterium glutamicum* showed that dimerization of SepF monomers is also possible by interaction of the lateral two α-helices forming a four-helix bundle. Moreover, it was observed that in this conformation two hydrophobic pockets are formed that enable interaction with FtsZ (16). In archaea, cell division systems are much less uniform than in bacteria. Three very different cell division mechanism have been identified in archaea: (i) a mechanism based on actin homologs, (ii) a mechanism based on homologs of the eukaryotic ESCRT-III system and (iii) a mechanism involving FtsZ (21). Studies on archaeal cell division have so far mainly focused on the crenarchaeal Cdv-system that is based on homologs of the ESCRT-III system also found in eukaryotes (22). Additionally, there are some studies on FtsZ-based euryarchaeal cell division, especially from Haloarchaea (23–29). Haloarchaea often contain multiple proteins belonging to the FtsZ/tubulin superfamily (30). A study in *Haloferax volcanii* showed that only two of the eight proteins from this superfamily are true FtsZ homologs. Both FtsZ homologs from *H. volcanii*, FtsZ1 and FtsZ2, were shown to be important for cell division and fulfill slightly different functions in the process (31). The remaining proteins form a distinct phylogenetic group named CetZ. Members of the CetZ group are not involved in cell division but in controlling the cell shape (31).

Interestingly, a bioinformatical approach identified putative archaeal SepF homologs in all FtsZ containing archaea, most of them belonging to the euryarchaeal superphylum (21). Putative archaeal SepF homologs from *Archaeglobus fulgidus* and *Pyrococcus furiosus* were used previously for crystallization studies (12). However, no functional studies on archaeal SepF homologs are available and it is currently unknown if these homologs are actually involved in cell division.

In this study, we used *H. volcanii* to characterize a putative archaeal SepF homolog. *Haloferax volcanii* is perfectly suited for the investigation of euryarchaeal cell biology due to the availability of an advanced genetic system (32) and functional fluorescent proteins (31), together with its easy cultivation conditions. Fluorescent microscopy of GFP-tagged SepF showed that it is localized at the cell center together with FtsZ1 and FtsZ2. Additionally, the gene is essential and characterization of its function was only possible upon depletion. SepF-depleted *H. volcanii* cells strongly increased in size over time and showed a cragged cell surface. On the other side, the presence of plasmids in the SepF depletion strain lead to filamentation of the cells upon SepF depletion. In contrast, overexpression of SepF had no effect on the growth and shape of *H. volcanii* cells. Moreover, localization of FtsZ1 and FtsZ2 in SepF-depleted cells might point to another so far unknown FtsZ1 membrane anchor. Indeed, an interaction of SepF with FtsZ2 but not with FtsZ1 was shown. In contrast to bacterial SepF, the archaeal SepF does not form polymers. In summary, we provide the first evidence that SepF is a key player in cell division in archaea using the FtsZ system.

## Results

### Localization of SepF in *Haloferax volcanii*

The first analysis indicating that FtsZ containing archaea might have SepF homologs was conducted by Makarova et. al in 2010 (21). In this study, archaeal clusters of orthologous genes (arCOGs) were identified with the same phyletic pattern as the arCOG containing the archaeal FtsZ homologs. arCOG02263 was identified as a cluster that seemed to contain archaeal SepF orthologs. This cluster is located on the main chromosome and contains the *hvo_0392* gene from *H. volcanii. hvo_0392* is not organized in an operon and adjacent genes encode proteins that are most likely not involved in cell division (Supplementary Figure 1A).

The protein encoded by *hvo_0392* is composed of 118 amino acids with a molecular weight of 12.6 kDA and an N-terminal membrane targeting sequence (MTS) (Supplementary Figure 1B). Comparison of the sequence of this protein with SepF sequences from selected bacteria and archaea showed a weak sequence similarity between the archaeal proteins and the SepF proteins from bacteria. Some of the residues of SepF, which were described to be important for interaction with FtsZ (12, 16), were conserved in both phyla (Supplementary Figure 1C). However, G109 of *B. subtilis* SepF which is important for oligomerization (12) and is conserved in other bacterial SepFs, is not present in archaeal SepF homologs (Supplementary Figure 1C).

To establish whether Hvo_0392 is involved in cell division, the gene was expressed with a C-terminal GFP-tag under the control of a tryptophan inducible promotor located on a plasmid (pSVA3942). Fluorescence microscopy showed localization of the protein in a ring-like structure at midcell (Figure 1A), at a similar location as observed for FtsZ1 in *H. volcanii* (35). At the resolution used, it was not possible to determine whether the rings were continuous or assembled from patches (Supplementary Figure 2). Localization of the tagged protein at the cell division plane in all observed cells (Figure 1B) strongly indicated that the protein encoded by the gene *hvo_0392* is involved in cell division and might have a similar function as bacterial SepF. Therefore, Hvo_0392 is referred to as SepF in this manuscript. SepF was localized at midcell in all imaged *H*.*volcanii* cells independent of the cell length, indicating that SepF is directly localized to the cell center after cell division. This was observed in both short cells (2 µm in length), that had just gone through cell division and elongated cells (6 µm in length), that were close to division (Figure 1B). In contrast, expression of SepF with an N-terminal GFP-tag showed diffuse localization throughout the cytoplasm (Figure 1C). Most likely this was caused by the GFP-tag blocking membrane binding of the N-terminal MTS of SepF (Figure 1D).

**Figure 1:**
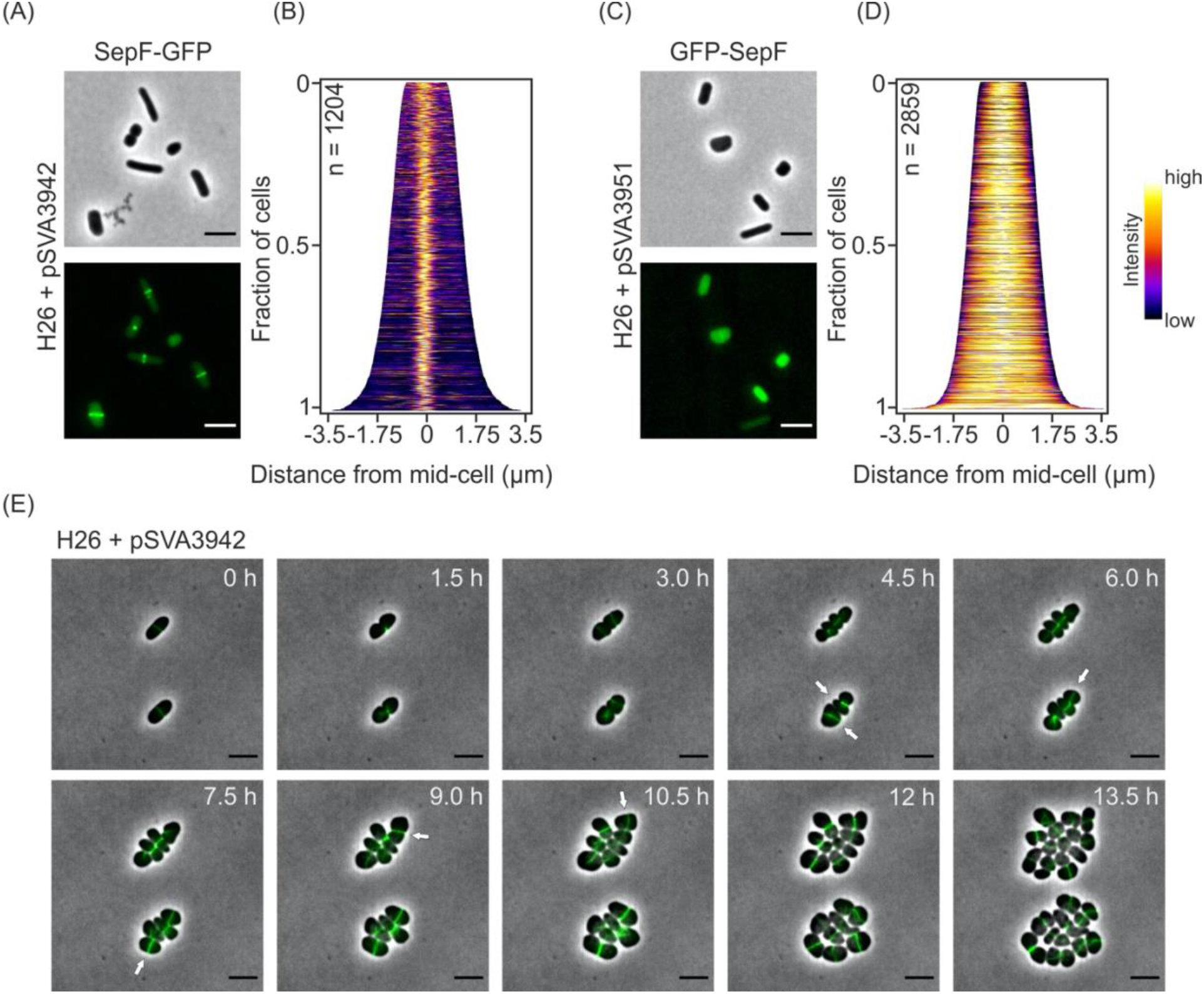
Localization and dynamic of SepF from *H. volcanii*. (A) Fluorescence microscopy of SepF-GFP expressing cells during early exponential growth. (B) Demographic analysis of the GFP signal in the imaged cells shows localization of SepF in the cell center independent of the cell length in all cells. (C) Fluorescence microscopy of GFP-SepF expressing cells during early exponential growth. (B) Demographic analysis of the GFP signal in the imaged cells shows diffuse signal of SepF throughout the cells independent of the cell length. Demographs were based on the GFP signals of cells imaged in three independent experiments. (E) Time-lapse microscopy of SepF-GFP expressing cells. Cells were grown on an agarose nutrition pad in a thermo-microscope set to 45 °C and imaged every 30 min. The selected pictures show constricting SepF rings during cell division. The new cell division plane in the daughter cells often occurs perpendicular to the old one (indicated by white arrows). Scale bars: 4 µm

**Figure 2:**
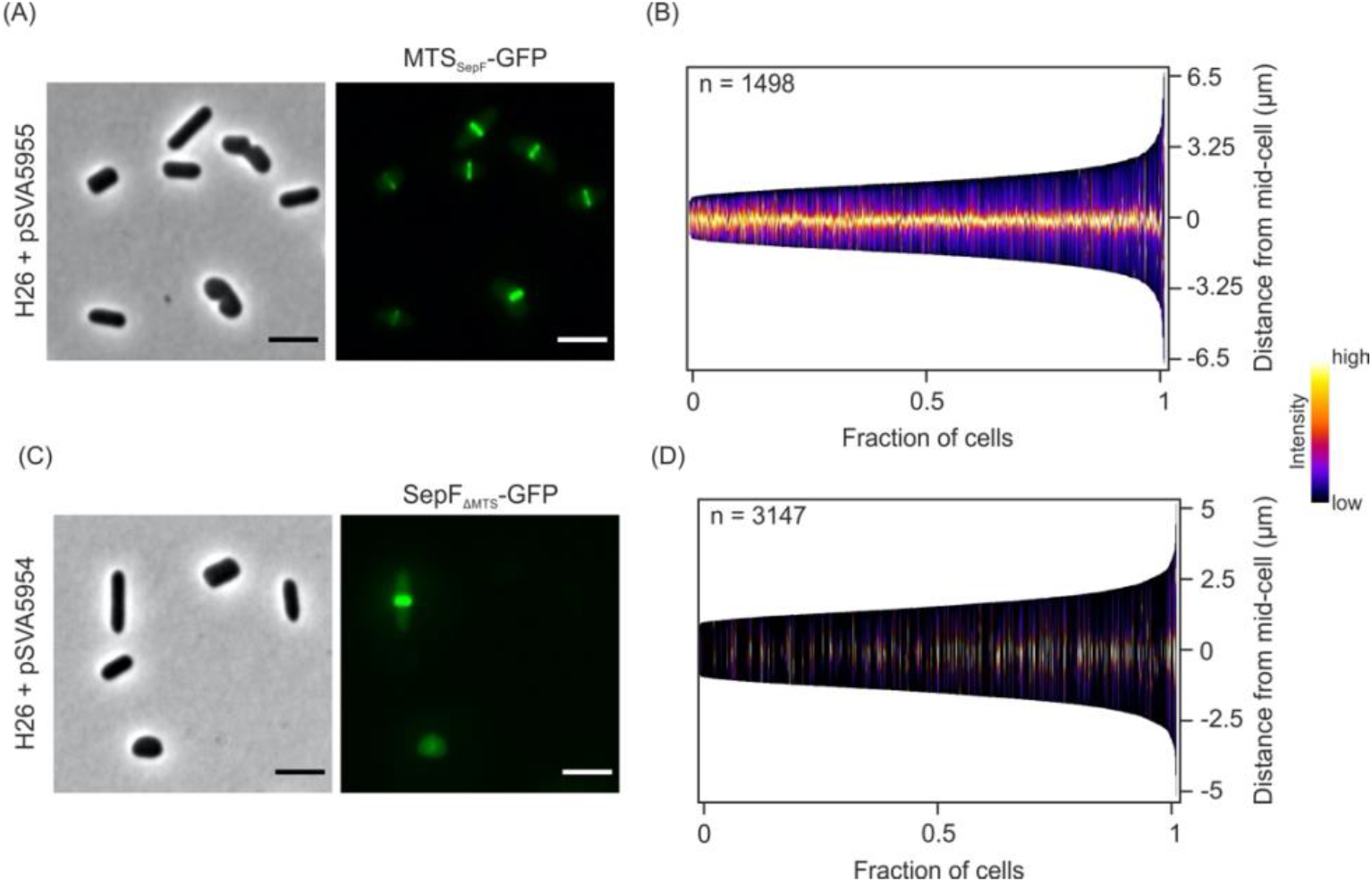
Localization of only the membrane targeting site of SepF and SepF without its membrane binding site. (A) Fluorescence microscopy of MTS_SepF_-GFP in *H. volcanii* cells during early exponential growth. Demographic analysis of the GFP signal in the imaged cells shows localization of the GFP-tagged membrane targeting site of SepF at midcell, independent of the cell length in all observed cells. (C) Fluorescence microscopy of SepF-GFP with the MTS of SepF deleted. (D) The construct was only expressed in 20 % of the cells. However, in these cells SepF without MTS was localized at midcell regardless of the cell length. Both demographs summarize the results of three independent experiments per construct. Scale bars: 4 µm.

To obtain insight into the dynamic behavior of the *H. volcanii* SepF homolog, time lapse microscopy was performed. *H. volcanii* cells expressing SepF-GFP were monitored for 13 h in a temperature-controlled microscope. During that time SepF rings were observed to constrict in the middle of the cells during cell division. Immediately after septum closure new SepF rings appeared at the future cell division plane in the daughter cells, often perpendicular to the old division plane (Figure 1E, Supplementary Video 1). To further investigate the influence of the MTS of SepF on the localization of the protein, the MTS sequence and SepF without its MTS were fused to GFP. Interestingly, the short peptide (MGIMSKILGGGG) forming the MTS of SepF was sufficient to efficiently localize GFP to the cell center in ring-like structures, like observed for the full length SepF above (Figure 2A, 2B). Moreover, SepF without its MTS was also shown to form ring-like structures at midcell possibly by interaction with endogenous SepF (Figure 2C). However, only 20 % of the observed cells (n = 3329) showed expression of the construct and its localization at midcell. In most of the cells, SepF-GFP without its MTS was not expressed (Figure 2D).

### Co-localization of SepF with FtsZ1 and FtsZ2

Since SepF-GFP was observed at midcell during cell division, it was tested whether it colocalizes with the tubulin homologs FtsZ1 and FtsZ2 at the midcell. Therefore, the respective genes were cloned in a double expression vector, expressing SepF with a C-terminal GFP-tag and FtsZ1 or FtsZ2 with a C-terminal mCherry-tag (Figure 3A, 3C). Both FtsZ1 (Figure 3B) and FtsZ2 (Figure 3D), localized together with SepF at midcell. However, while the signals of SepF and FtsZ2 strongly overlapped, the FtsZ1 signal was slightly broader than the SepF signal. These results show that SepF localizes like FtsZ1 and FtsZ2 at midcell during cell division, indicating that SepF might be a part of the divisome of *H. volcanii*.

**Figure 3:**
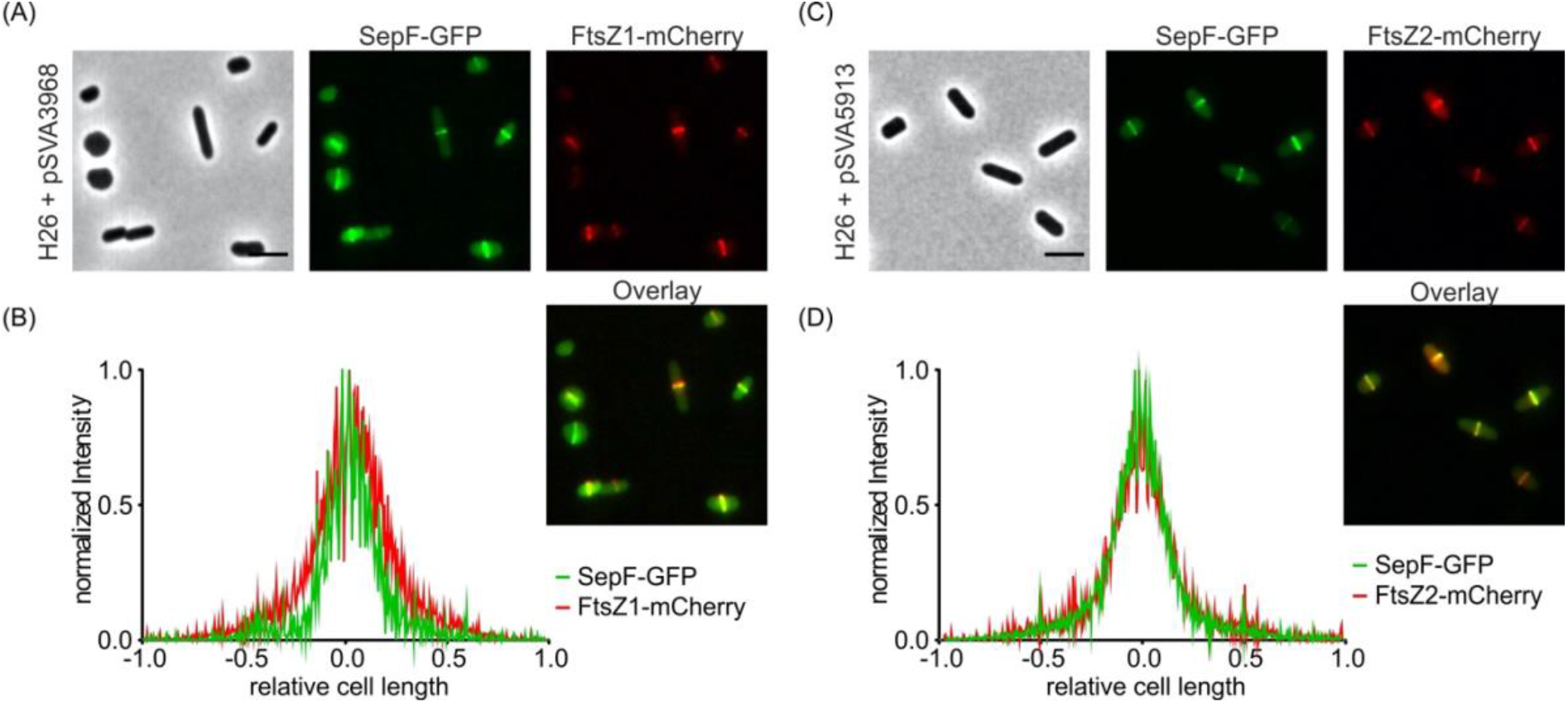
Co-localization of SepF with either FtsZ1 or FtsZ2. (A) Fluorescence microscopy of *H. volcanii* cells expressing SepF-GFP and FtsZ1-mCherry in early exponential phase. (B) Intensity profile of the normalized fluorescence signal from the SepF-GFP (green) and the FtsZ1-mCherry (red) construct relative to the cell length.Fluorescence microscopy of *H. volcanii* cell expressing SepF-GFP and FtsZ2-mCherry in early exponential phase. (D) Intensity profile of the normalized fluorescence signal from the SepF-GFP (green) and the FtsZ2-mCherry (red) construct relative to the cell length. The experiment was repeated at least three times per construct with > 1000 cells analyzed. Scale bars: 4 µm.

### SepF is essential in *Haloferax volcanii*

To obtain insight into the cellular function of archaeal SepF several attempts to delete *sepF* were made. However, we were not able to generate a knock-out mutant of *sepF* suggesting an essential function for SepF. To test this hypothesis, the native promotor of the gene was exchanged by a tryptophan inducible promotor resulting in the strain HTQ239. The HTQ239 strain showed normal growth when grown in selective medium complemented with 0.25 mM tryptophan (Figure 4A). After dilution into selective medium without tryptophan the culture stopped growing whereas the control strain H26 grew unaffected (Figure 4B), demonstrating that SepF is essential *H. volcanii*. To further study the effects of SepF depletion, time lapse microscopy of HTQ239 cells on an agarose nutrition pad without tryptophan were performed. This showed that the cell size strongly increased over time (Figure 4C). The cells failed to establish a cell division plane, resulting in bloated cells, which is a phenotype similar that that observed when an inactive form of FtsZ1 (D250A) is expressed in *H. volcanii* (31). The increase of the cell size also explains why the optical density of the culture of the SepF depletion strain still slowly increased. The severe cell division defect after SepF depletion and the strong increase of the cell size implies an important role of SepF in the division process. Besides functioning as a possible membrane anchor for FtsZ1 or FtsZ2, SepF might also be involved in the regulation of cell division.

**Figure 4:**
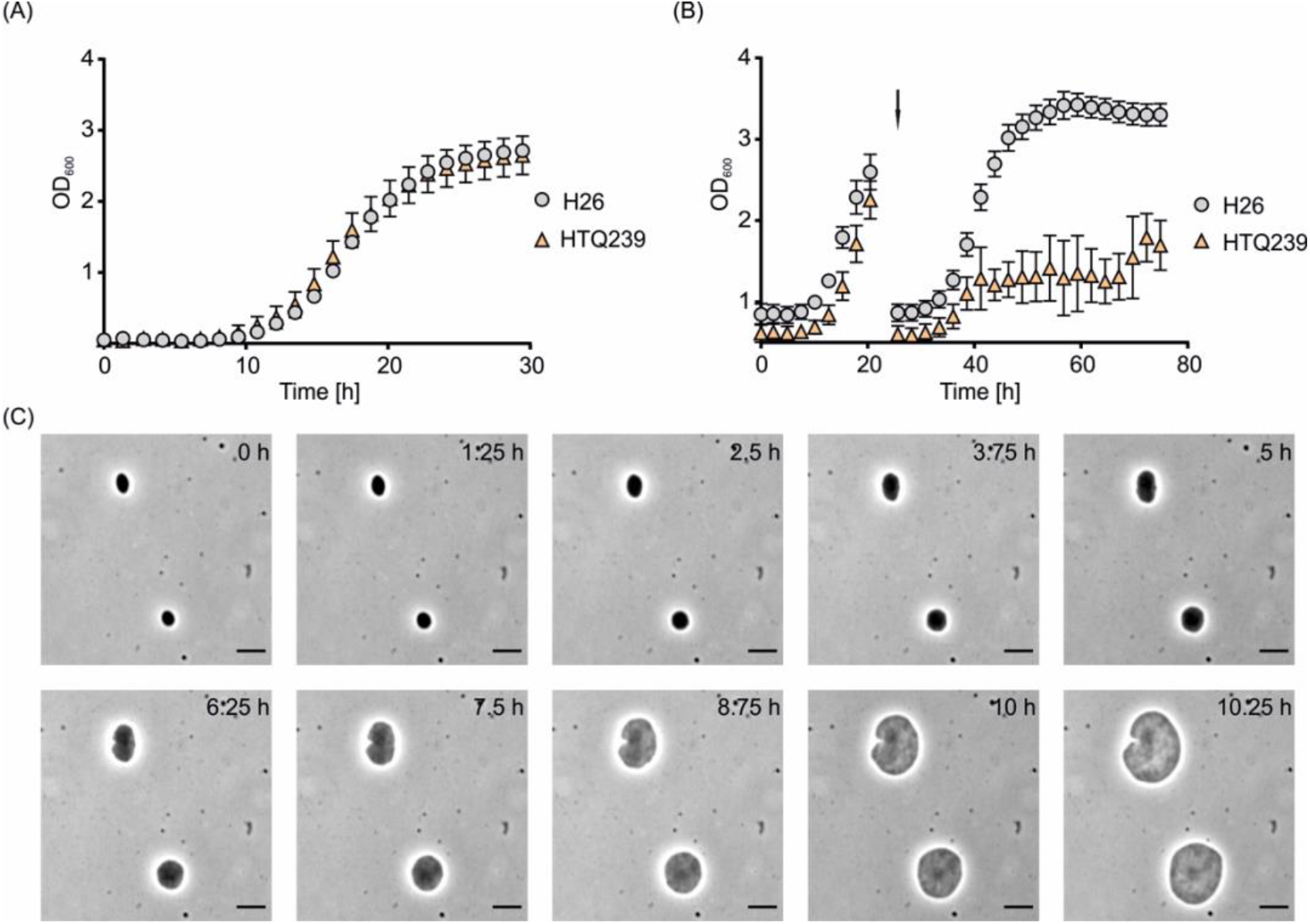
Growth of H26 and HTQ239 in the presence and absence of tryptophan. (A) Growth curve of H26 (grey) and HTQ239 (orange) grown in CAB-medium supplemented with tryptophan and uracil. (B) Growth curve of H26 (grey) and HTQ239 (orange) after dilution in CAB-medium without tryptophan (indicated by the arrow). After the transfer, HTQ239 stopped growing while H26 grew unaffected. The data for this growth curves were acquired from four independent cultures per strain. (C) Time lapse microscopy of HTQ239 cells in the absence of tryptophan: Cells were grown on an agarose nutrition pad (without tryptophan) in a thermo-microscope set to 45 °C and imaged every 15 min. During imaging the cells strongly increased in size while they were not able to execute cell division. Exemplarily cells every 1.25 h after depletion are shown. Scale bars: 4 µm

### Overexpression of *sepF* in *Haloferax volcanii* has no effect on growth and cell shape

Several studies in bacteria reported that overexpression of *sepF* hampers cell division (17, 18). To study whether overproduction of the archaeal SepF also leads to impaired cell division, *sepF* was cloned in plasmid pTA1992 (resulting in pSVA5960) that allowed constitutive SepF expression. The investigated strains were first grown in medium with tryptophan and diluted later into selective medium without tryptophan while the OD_600_ was constantly measured (Figure 5).

**Figure 5:**
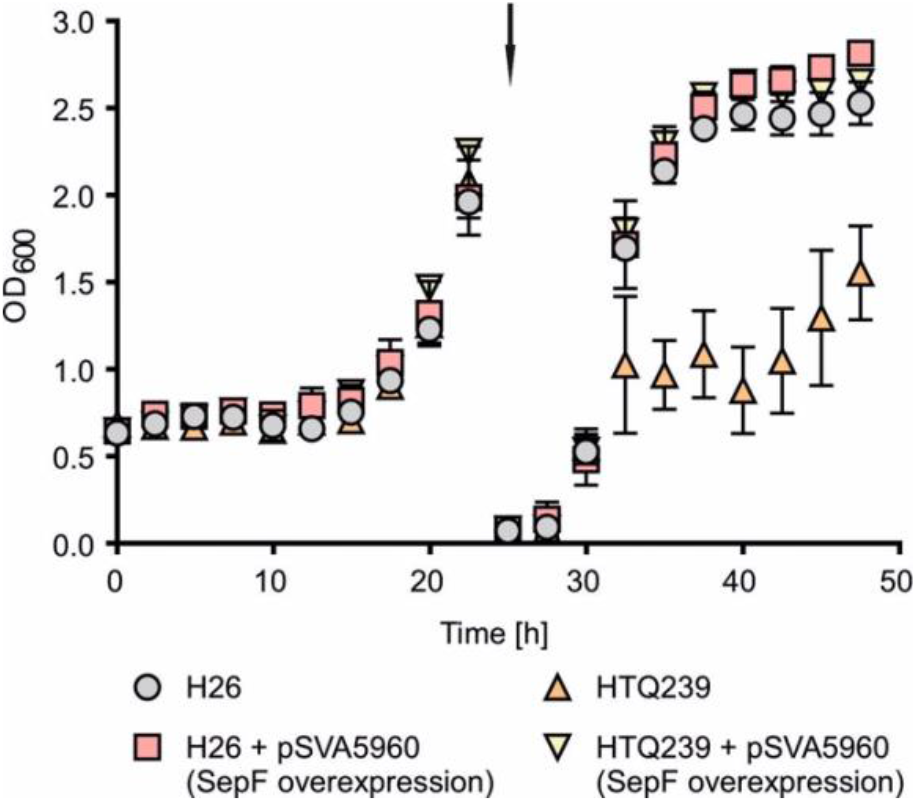
Growth curve of H26 and HTQ239 overexpressing SepF in the presence and absence of tryptophan. Growth curve of H26 (grey), H26 + pSVA5960 (red), HTQ239 (orange) and HTQ239 + pSVA5960 (yellow) grown in CAB-medium supplemented with tryptophan and uracil. All four strains grew the same before dilution into CAB-medium without tryptophan (indicated by the arrow). The data for this growth curve were acquired from three independent cultures per strain.

Like observed before, the SepF depletion strain stopped growing after dilution into medium without tryptophan. Overexpression of *sepF* in the wild-type strain had no effect on the growth as compared to H26 without an overexpression plasmid. Both strains grew in a similar manner in medium with and without tryptophan. Moreover, overexpression of *sepF* in the *sepF* depletion strain HTQ239 complemented the growth defect caused by the SepF depletion (Figure 5). To further analyze the effect of overexpression of *sepF* in strains H26 and HTQ239 cells were imaged every three hours after dilution into medium without tryptophan (Figure 6A) and the cell length (Figure 6B) and area (Figure 6C) were determined. Before dilution into medium without tryptophan all strains were rod-shaped with a length ranging from 3-4 µm (t = 0). During the next nine hours cells of strain H26, H26 + pSVA5960 and HTQ239 + pSVA5960 slowly changed their cell shape from a rod- to a plate-shaped form and the cell length was reduced to 3 µm as expected for normal growth of *H. volcanii* (36). In contrast, the length of the depletion strain strongly changed within nine hours. Already after three hours in medium without tryptophan HTQ239 cells showed an increased length compared to H26. Six hours after SepF depletion the cells had two times the length of H26 at the same time point and three hours later cells had three times the length of H26 (Figure 6B). Moreover, the cell shape of the depletion strain not only changed in length but also in width, resulting in a strongly increased cell size within nine hours. After 3, 6 and 9 hours in medium without tryptophan, HTQ239 cells showed 2, 5 and 6 times larger cell area then H26 cells (Figure 6C). After 24 hours in medium without tryptophan, H26, H26 + pSVA5960 and HTQ239 + pSVA5960 were plate-shaped with an average length of 1.8 µm, the typical cell morphology for *H. volcanii* cells in the stationary phase. The cells of HTQ239 without any plasmid were partially hugely bloated or disrupted and most of the cells had a cragged cell surface. Due to the different grey values the HTQ239 cells had in the acquired images at the last time point it was no longer possible to measure the size with the image analysis software.

**Figure 6:**
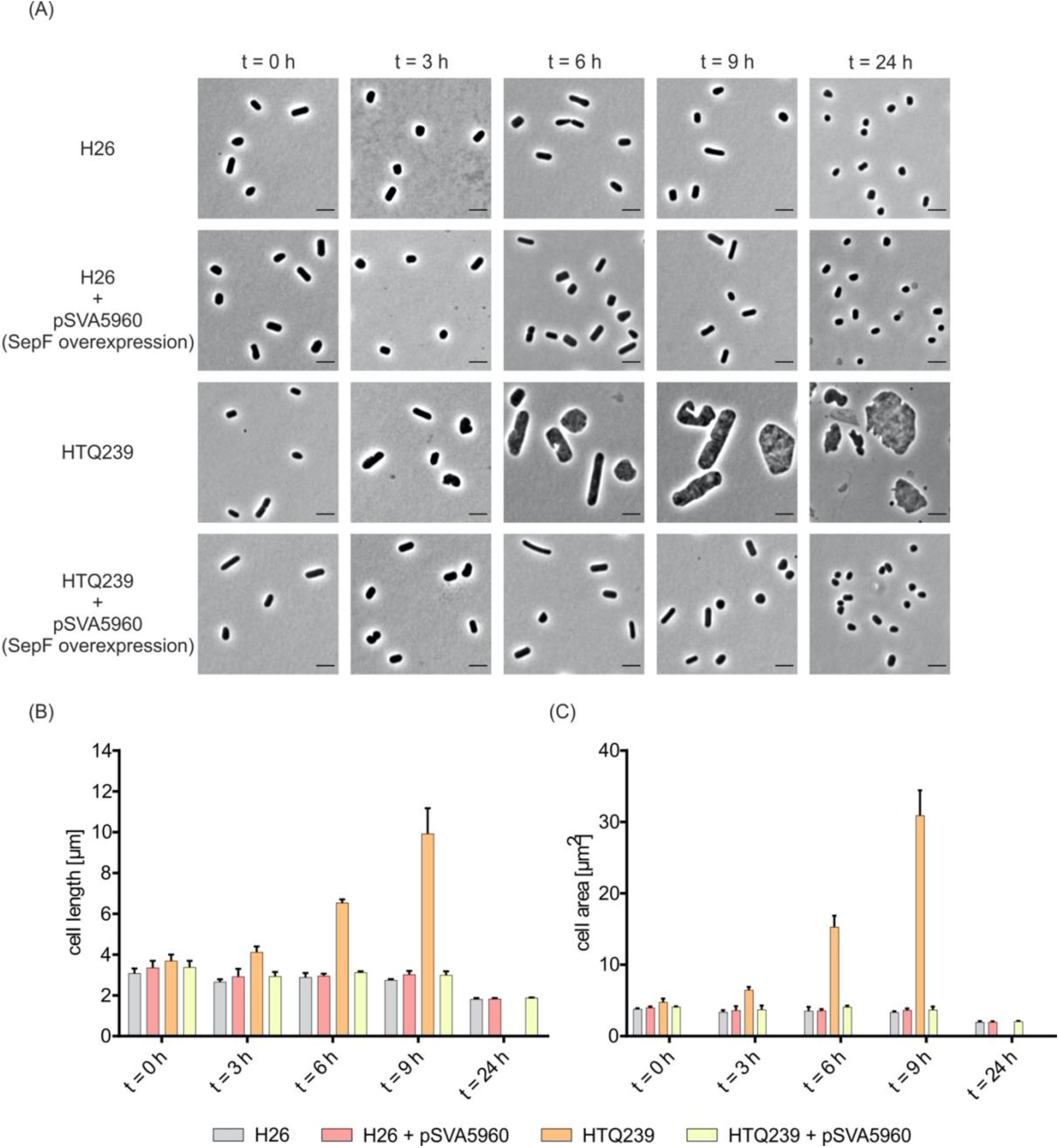
Cell shape analysis of H26 and HTQ239 during *sepF* overexpression in the absence of tryptophan. (A) Microscopy of H26 and HTQ239 cells with or without SepF overexpression plasmid pSVA5960. Cells were imaged in 3 h steps after SepF depletion for 9 h and one last time 24 h after depletion. Scale bar: 4 µm. (B) Mean cell length of strain H26 (grey), H26 + pSVA5960 (red), HTQ239 (orange) and HTQ239 + pSVA5960 (yellow) at different time points after SepF depletion. (C) Mean cell area at different time points after SepF depletion. Cell shape parameters were obtained from three independent experiments including the cell size of > 1000 cells per strain and time point.

### Localization of FtsZ1-GFP and FtsZ2-GFP in SepF-depleted *H. volcanii* cells

In order to study the effect of the SepF depletion on the localization of FtsZ1 and FtsZ2, strain HTQ239 was transformed with either a plasmid expressing *ftsZ1-gfp* (pSVA5910) or a plasmid expressing *ftsZ2-gfp (*pSVA5956) under the control of their own promotors. Expression of either *ftsZ1-gfp* or *ftsZ2-gfp* in the HTQ239 strain resulted in filamentation of the cells after SepF depletion (Supplementary Figure 3A). Importantly, this filamentation phenotype is caused by the presence of the plasmid in the *sepF* deletion strain. Transformation of the backbone plasmid pTA1392, not expressing either *ftsZ1-gfp* or *ftsZ2-gfp* in HTQ239 and subsequent SepF depletion, resulted in the filamentation phenotype, while H26 transformed with pTA1392 maintained a normal cell shape (Supplementary Figure 3). Why the presence of the plasmid in the deletion strain causes the filamentation phenotype is currently unknown, but It has been observed previously that the presence of a plasmid in *H. volcanii* strains with deletions which are linked to cell shape or division leads to a higher amount of rod-shaped cells compared to cells without plasmid at the same growth stage (37, 38). Like before, cells were imaged every three hours after SepF depletion. Over time, the cell length of both strains strongly increased compared to the wild type. Nine hours after dilution into medium without tryptophan HTQ239 expressing *ftsZ1-gfp* had an average cell length of 12.6 µm and HTQ239 expressing *ftsZ2-gfp* an average length of 13.8 µm, almost five times longer than H26 at the same time. Twenty-four hours after SepF depletion cells were still filamentous but without further increase of the cell length compared to the cells imaged fifteen hours before (Supplementary Figure 3B). Interestingly, depletion of SepF had a different effect on FtsZ1 localization compared to FtsZ2 localization. Before dilution into medium without tryptophan FtsZ1 localized to midcell in strain HTQ239 in almost all observed cells (Figure 7A). Three hours after depletion the cells started to increase in size but most of the cells still had one FtsZ1 ring. Only cells with a length of more than 6 µm started to assemble additional rings. With increasing cell length (t = 3 and t = 9) additional FtsZ1 rings were observed in the elongated cells. After 24 hours of SepF depletion, cells with up to five FtsZ1 rings, evenly distributed over the cell body, were detected. Due to the different numbers of FtsZ1 rings in the cells, the demographs appear rather noisy. However, the demographs from nine and 24 hours after dilution into medium without tryptophan showed that one FtsZ1 ring was always located close to one of the two cell poles (Figure 7A). Before SepF depletion HTQ239 cells expressing *ftsZ2-gfp* showed one FtsZ2 ring per cell located in the cell center (Figure 7B).

**Figure 7:**
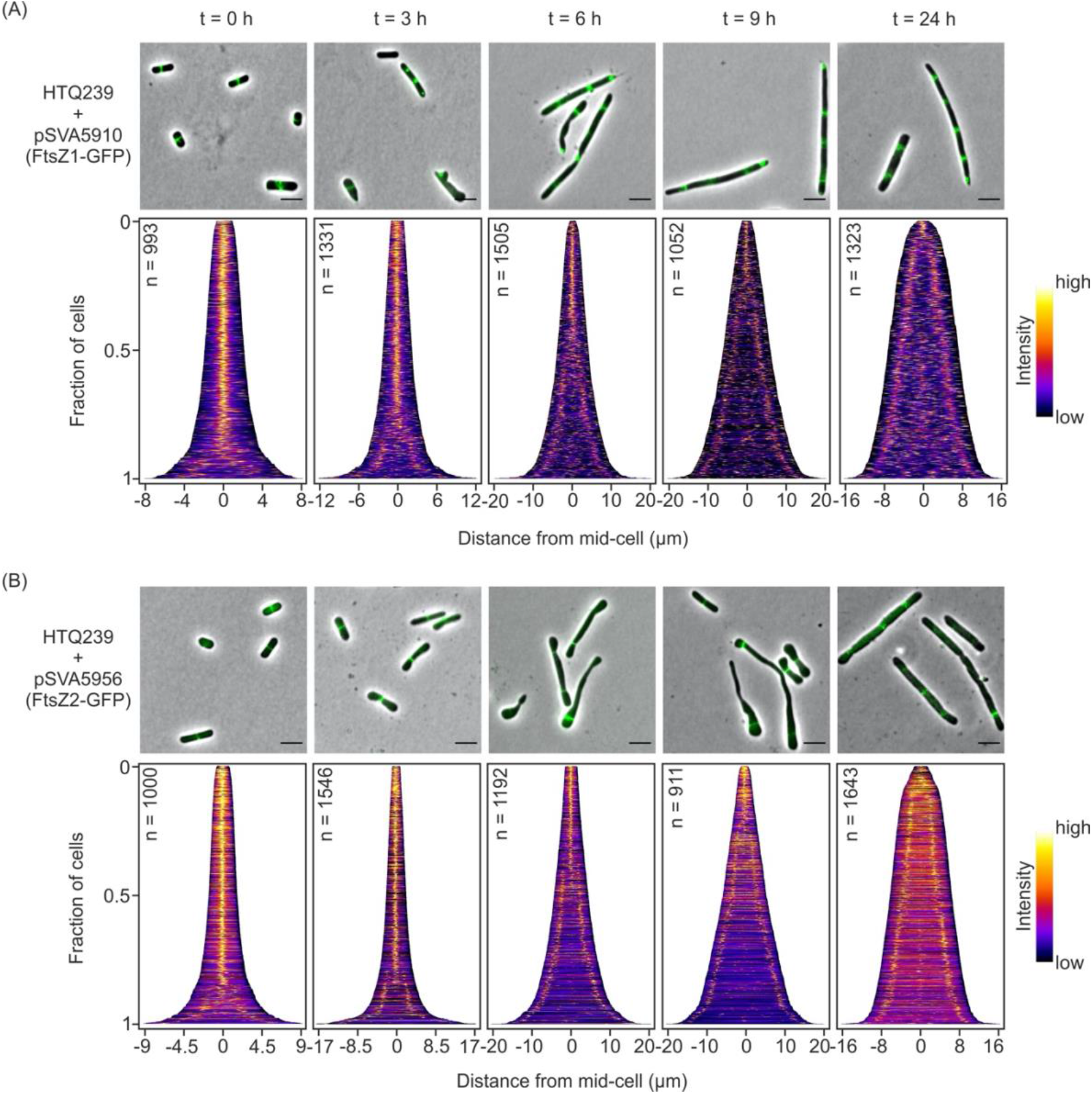
Localization of FtsZ1 and FtsZ2 in SepF-depleted cells. (A) Fluorescence microscopy of HTQ239 cells additionally expressing *ftsZ1*-*gfp* under the control of its native promotor at different time points (0, 3, 6, 9 and 24 h) after sepF depletion. Over time the cells became filamentous and the number of FtsZ1 rings increased up to five rings per cell. (B) Demographic analysis of the FtsZ1-GFP signal observed in HT239 at the indicated time points after SepF depletion. The number of cells analyzed per time point is indicated in the respective demograph. (C) Localization of FtsZ2-GFP by fluorescence microscopy in HTQ239 at different time points after SepF depletion. Cells became filamentous over time and the number of FtsZ2 rings partially increased up to three rings per cell. (D) Demographic analysis of the FtsZ2-GFP signal imaged in the SepF-depleted HTQ239 cells at different time points. The number of cells analyzed per time point is indicated in the respective demograph. The demographs summarize the results from three independent experiments per strain. Scale bars: 4 µm

became filamentous as well. Cells longer than 10 µm had two FtsZ2 rings while cells shorter than 10 µm had only one FtsZ2 ring. In contrast to the number of FtsZ1 rings observed in HTQ239 cells, a maximum of three FtsZ2 rings were observed per cell. Moreover, in elongated cells (t = 6–24 h, > 10 µm) with only one FtsZ2 ring the ring was always located close to one cell pole. In cells with two FtsZ2 rings each ring was located near one of the two poles. The third FtsZ2 ring in cells that had three rings was located at midcell between the other two FtsZ2 rings. As observed for FtsZ1, the FtsZ2 rings appeared in a distinct distance along the cell body (Figure 7B). Since SepF was shown to directly interact with FtsZ in bacteria a decrease of FtsZ1 and FtsZ2 rings after SepF depletion was expected for *H. volcanii*. Instead, the number increased to up to three rings per cell for FtsZ2 and up to five rings for FtsZ1. It is possible that the remaining SepF after depletion was sufficient to anchor up to three FtsZ2 rings to the membrane. However, the high number of additional FtsZ1 rings in SepF-depleted cells might point to another, yet unknown, FtsZ1 membrane anchor instead of SepF.

### SepF from *Haloferax volcanii* interacts with FtsZ2 but not with FtsZ1

To further investigate whether SepF directly interacts with FtsZ1 or FtsZ2, pulldowns from the cytosolic or membrane fraction of cross-linked SepF-HA expressing cells (HTQ236) were performed using magnetic beads coated with antibodies against the HA-tag. The pulldown fractions were subsequently analyzed via Western blot analysis using specific antibodies against either FtsZ1 or FtsZ2, and by mass spectrometry. Mass spectrometry could not identify any peptides of FtsZ1 in the elution fractions of either the control sample from H26 or the SepF-HA expressing strain HTQ236 (Supplementary Table S4), whereas SepF was present in high amounts in the pulldown fractions from the HTQ236 strain. The same result was obtained using Western Blot analysis with α-FtsZ1 and α-FtsZ2 antibodies. Whereas only an unspecific signal was detected for the α-FtsZ1 antibody in all pulldown fractions (Figure 8A), a very clear signal indicating the presence of FtsZ2 specifically in the pulldown fraction from the membranes from the HTQ236 strain (Figure 8B). These pulldown assays thus demonstrated a direct interaction between SepF and FtsZ2.

**Figure 8:**
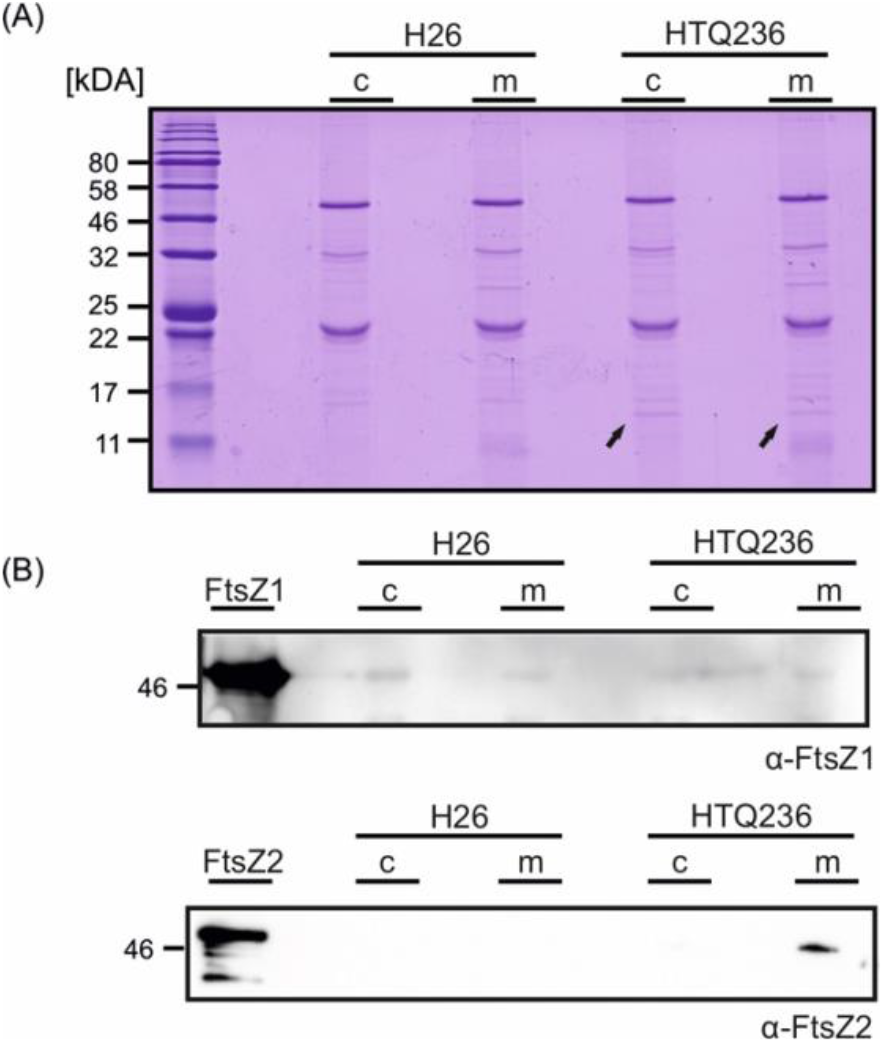
Immunoprecipitation of SepF-HA. Cytosolic (c) or membrane (m) fractions used for immunoprecipitation from H26 or HTQ236 were loaded on 15 % SDS-gels (A) and analyzed via Western blots using antibodies either against FtsZ1 or FtsZ2 (B). The HA-tagged SepF was visible on the Coomassie stained gels in all pulldown fractions of strain HTQ236 (indicated by the black arrows).

### *H. volcanii* SepF does not form oligomers *in vitro*

Bacterial SepF was reported to form oligomers by lateral interaction between SepF dimers (12). Since G109 of *B. subtillis* SepF, which in bacteria is important for this interaction, is not conserved in archaeal SepF homologs, it was tested whether *H. volcanii* SepF could also form oligomers. *H. volcanii* SepF was expressed in *E. coli* and purified by nickel affinity-chromatography (Supplementary Figure 4AB) and size exclusion chromatography experiments were performed. SepF eluted at a position corresponding to a molecular weight of ∼ 30 kDa, corresponding to a dimeric protein and no protein eluted at elution volumes corresponding to higher oligomeric complexes (Figure 9A). The eluted protein was analyzed via Western blot using antibodies against the His-tag to verify that the eluted protein is SepF (Figure 9B). Thus *H. volcanii* SepF forms a dimer and no indication of oligomerization could be found *in vitro*.

**Figure 9:**
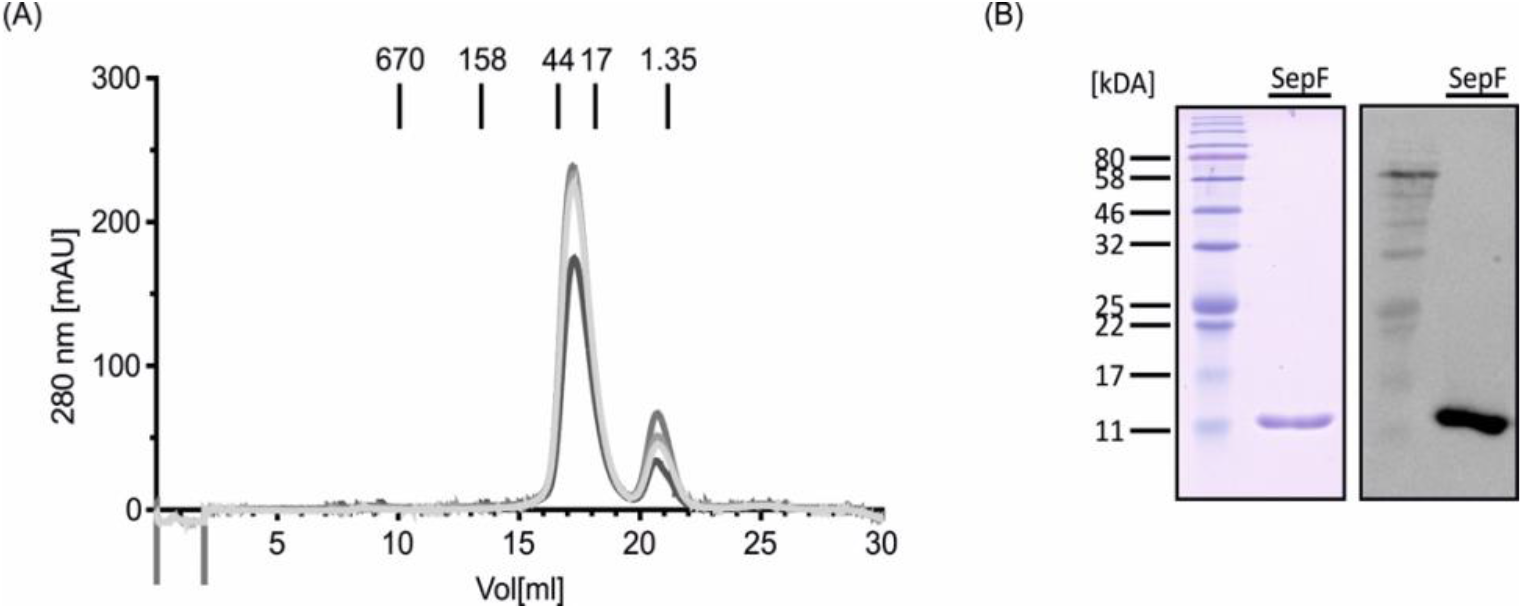
Size exclusion chromatography with SepF. (A) Purified SepF was loaded on a Superdex 200 10/300 GL column. Absorption at 280 nm at the different elution volumes for 4 independent experiments are shown. Elution volumes of protein size markers are indicated in kDA. (B) SDS-gel and Western blot with anti-His antibodies of SepF after SEC.

## Discussion

SepF is an important protein involved in the cell division process of several bacterial species, including Cyanobacteria, Firmicutes and Actinobacteria, functioning as an alternative membrane anchor for FtsZ (12, 13). Genes encoding SepF homologs are also found in FtsZ containing archaea (21). Studies of archaeal SepF homologs were until now limited to the elucidation of the crystal structure of the putative SepF proteins from *A. fulgidus* and *P. furiosus* (12). Very recently, the crystal structure of SepF of *Methanobrevibacter smithii* in its apo-form and in complex with the C-terminal domain of FtsZ was solved (Pende N., Sogues A., et al., personal communication) demonstrating that SepF also directly interacts with FtsZ in archaea. We present here, the first functional study of an archaeal SepF protein and show that *H. volcanii* SepF is an essential component of the archaeal cell division machinery.

Fluorescence microscopy of SepF-GFP in *H. volcanii* showed localization in a ring-like structure at the cell division plane in the center of the cell, similar to the localization of bacterial SepF (18), which was also observed by immunofluorescence in the archaeon *M. smithii* (Pende N., Sogues A., et al., personal communication). Similar to the SepFs from bacteria, SepF from *H. volcanii* contains an N-terminal membrane targeting site (MTS). An N-terminal GFP fusion resulting in a diffused localized GFP-SepF, suggesting the GFP prevented the membrane localization of SepF. Interestingly, the short MTS sequence fused to GFP, efficiently localized the fluorescent protein to midcell. Hence, besides anchoring SepF to the cell membrane, the MTS of archaeal SepF also is important for midcell localization. Indeed, time lapse microscopy of SepF-GFP producing cells showed that SepF is localized at the future site of cell division immediately after closure of the septum, making SepF one of the first cell division proteins to arrive at the new septum. It is known that the future cell division plane in *H. volcanii* is assembled early during the cell cycle because FtsZ1 and FtsZ2 arrive shortly after the end of the previous round of cell division site (31, 38). However, the assembly hierarchy of SepF, FtsZ1 and FtsZ2 is unknown so far. We show here that FtsZ2 interacts with SepF, suggesting that SepF functions as a membrane anchor which brings FtsZ2 to the division site. In contrast, we were not able to show direct interaction between SepF and FtsZ1. This leads to the assumption that SepF is an exclusive membrane anchor for FtsZ2 and that there is another, likely non-canonical, membrane anchor for FtsZ1 present in archaea that involve two FtsZ homologs for cell division. Moreover, FtsZ2 localization was reported to highly depend on the presence of FtsZ1 (38). This indicates that FtsZ1 might be the first cell division protein that is placed at the future site of cell division. The mechanism for spatial and temporal organization of Z-ring positioning in archaea is still unknown. Homologs of the MinD protein are present in *H. volcanii* but were recently shown not to be involved in Z-ring positioning, in contrast to the bacterial MinD protein (39). Our SepF depletion experiment showed that SepF had no negative effect on FtsZ1 ring formation as the number of additional FtsZ1 rings increased to up to five rings during late stages of depletion. However, only a maximum of three FtsZ2 rings were seen during SepF depletion. Possibly their formation was triggered by SepF remnants from before depletion. Interestingly, the presence of a plasmid in SepF depletion strain HTQ239 resulted in filamentous cells. In these filamentous *H. volcanii* cells we observed that the additional FtsZ1 rings were formed equidistanced. This hints at an oscillating negative regulator for FtsZ1-ring positioning in Euryarchaeota, analogous to the bacterial Min-system. From bacteria it is known that the cell length directly influences the oscillation pattern of the Min-system. In cells with a normal length pole-to-pole oscillation of the Min-system is observed that changes to a multi-node standing wave in filamentous cells (40–42). This alteration in the oscillation of the Min-system results in regions with minimal concentrations of the negative Z-ring assembly regulator MinC, leading to the formation of equally distributed FtsZ rings (42), similar to the FtsZ1 rings we observed in filamentous *H. volcanii* cells. This observation implies that the localization of the cell division plane in Euryarchaeota is determined by FtsZ1 positioning and spatially regulated by a so far unknown oscillating mechanism. Investigation on SepF from *Mycobacterium tuberculosis* showed that the positioning of SepF is dependent on FtsZ (18). This was recently confirmed as interaction of SepF with FtsZ was crucial for its membrane binding in *Corynebacterium glutamicum* (16). Possibly, FtsZ1 triggers membrane interaction of SepF in *H. volcanii* by a transient interaction and in turn SepF recruits FtsZ2 to the site of cell division, forming the early divisome.

From bacteria there is evidence that the SepF protein is also involved in later steps of cell division, indicating that SepF also has a regulatory role during cytokinesis (13, 16–18, 43–46). Therefore, SepF needs to be tightly regulated as changes of native SepF levels leads to severe cell division defects. Especially overexpression of SepF was shown to completely block cell division in different bacteria (16–18). Interestingly, additional overproduction of FtsZ mitigated the effect of SepF overexpression, implicating that the SepF/FtsZ ratio and the reduction of freely diffusing SepF is crucial for correct cell division in bacteria (17). In contrast, no negative effect was observed in SepF overexpressing *H. volcanii* cells and, moreover, SepF overexpression rescued the SepF-depleted cells. A recent study stated that *sepF* and *ftsZ2* in *H. volcanii* are under the control of the same transcription factor and thus concluded that both protein levels are co-regulated (Vogel et al., Iain Duggin personal communication). The exact intracellular concentrations of SepF and FtsZ2 are unknown but the ratio of both proteins apparently does not need to be as tightly regulated as in bacteria since SepF overproduction and thereby the increased concentration of free archaeal SepF does not block cell division like observed for bacteria (17). Nevertheless, we assume that SepF has also a regulatory role in *H. volcanii* as during SepF depletion constriction of the formed FtsZ1 and FtsZ2 rings was blocked and no septum closure had been observed. The force for septum closure in bacteria is generated by incorporation of new cell wall material at the septum (3–5) and free SepF in *B. subtilis* was reported to lead to delocalization of proteins of the septal peptidoglycan (PG) synthesis machinery (17). Moreover, in *M. tuberculosis* SepF was reported to interact with MurG, a protein involved in maturation of the peptidoglycan precursor Lipid-II (45). In another actinobacterium, *Corynebacterium glutamicum*, no septal PG incorporation has been observed after SepF depletion (16).

A high variety of cell enevlopes is found in archaeal species. For example the cell envelope of *H. volcanii* only consist of a proteinaceous lattice called the S-Layer while *M. smithii* and other Methanobacteriales have a cell wall polymer named pseudomurein that is similar to the bacterial peptidoglycan (47, 48). During SepF depletion in *H. volcanii* we observed a strong increase in the cell size, most likely the result of uncoordinated incorporation of new cell wall material all over the cell body. This suggests that the archaeal SepF is also involved in the coordination of new cell wall material incorporation. This indicates that SepF localization at the septum is crucial for positioning the cell wall synthesis machinery to the site of cell division in archaea. Indeed, previous studies reported that in several archaeal species new S-Layer is incorporated at the cell center (49–51). Additionally, in *H. volcanii* the authors observed that archaeosortase A (ArtA), a protein that processes and links several proteins like the S-Layer glycoprotein to a lipid moiety called archaetidylethanolamine, is localized at midcell during cell division (51). Moreover, two proteins that are assumed to be involved in archaetidylethanolamine synthesis, PssA and PssD, were observed at the cell center as well (51). However, as we did not detect these proteins in our immunoprecipitations experiment, we assume that SepF does not directly interact with them and their possible delocalization is a downstream effect of SepF depletion. Furthermore, as SepF shows the same localization pattern in archaea with different cell envelopes, *H. volcanii* (our study) and *M. smithii* (Pende N., Sogues A., et al., personal communication), it is likely that archaeal SepF is capable to coordinate completely different machineries for septal cell envelope incorporation in archaea with different cell sheaths that have a FtsZ based cell division machinery.

It is possible that SepF functions as a linker between early and late cell division proteins and switches the divisome from an “off” to an “on” state, similar to FtsA in bacteria. FtsA signaling during cell division is assumed to be achieved by transition from a polymeric to a monomeric state (52, 53). However, size exclusion experiments with the archaeal SepF suggest only dimer formation and moreover the conserved glycine 109 in bacterial SepF, important for polymerization (12), is not present in the archaeal SepF protein. Moreover, also purified SepF from *M. smithii* did not form higher ordered structures as well and was only found in a dimeric conformation (Pende N., Sogues A., et al., personal communication).This indicates that SepF in archaea is a dimer that does not form oligomers. Polymers of bacterial SepF were capable to remodel membranes, possibly initiating membrane invagination at the septum (12, 16). Unexpectedly, the archaeal SepF was shown to remodel small unilamellar vesicles without the formation of higher ordered SepF structures (Pende N., Sogues A., et al., personal communication).

Based on our study on the archaeal SepF protein and the previously reported study on both FtsZ homologs in *H. volcanii* (38), we propose the following model for divisome formation in Euryarchaeota: Directly after cell division, FtsZ1 is localized to the future site of cell division by a so far unidentified, oscillating, negative regulator. Subsequently, FtsZ1 protofilaments are attached to the membrane by a so far unknown non-canonical membrane anchor forming the Z1-ring. SepF is recruited to the cell center by a transient interaction with FtsZ1, possibly via the MTS of SepF. SepF dimers in turn tether FtsZ2 filaments to the membrane resulting in the formation of the second Z-ring and the completion of the early divisome. The formed Z2-ring then provides a scaffold for further downstream cell division proteins including ArtA, PssA and PssD. Incorporation of new S-Layer at the cell center would then lead to septum closure and separation of the newborn daughter cells (Figure 10). We are aware that this is a simplified model as we expect many more proteins to be part of the euryarchaeal divisome that yet have to be identified. Moreover, it is likely that only the proteins of the early divisome like FtsZ1, FtsZ2 and SepF are conserved in Euryarchaeota because the cell envelopes differ strongly amongst euryarchaeal species (47, 54).Hence, it can be assumed that proteins for late steps in cell division are not conserved amongst Euryarchaeota as also the machineries for cell wall material incorporation at the septum need to be adapted to the different types of cell envelopes.

**Figure 10:**
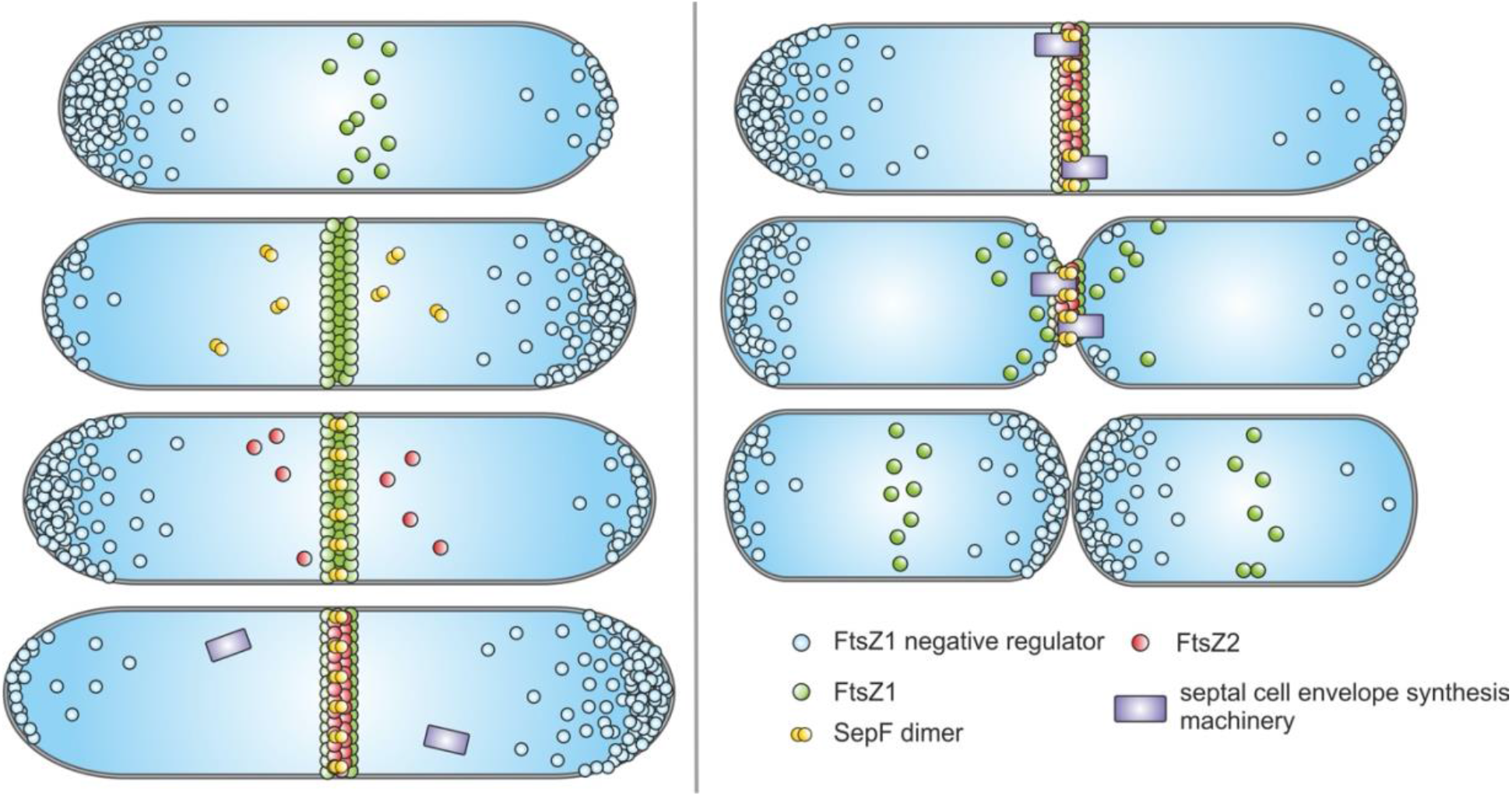
Model of cell division in *H. volcanii*. Left: Directly after the last round of cell division FtsZ1 (green) is positioned at the future site of cell division, by a so far unidentified oscillating negative regulator (blue), forming the FtsZ1 ring. Next, SepF dimers (yellow) are localized in a FtsZ1 dependent manner to the site of cell division and SepF recruits and anchors FtsZ2 (red) to the septum, resulting in the formation of the FtsZ2 ring. Right: After assembly of the early divisome proteins for septal cell envelope synthesis (purple) are recruited to midcell. Possibly, SepF triggers incorporation of new cell wall material that leads to constriction of the septum and segregation of the newborn daughter cells. From the septum released FtsZ1 proteins then directly localize to the future site of cell division.

In conclusion, the predicted archaeal SepF protein is an essential part of the euryarchaeal divisome that not only provides a membrane anchor for FstZ2 but is also important for the regulation of the machinery for new cell wall material incorporation at the cell center.

## Material and Methods

Unless otherwise stated all chemicals and reagents were obtained either from Carl Roth or Sigma-Aldrich.

### Strains and growth media

*Haloferax volcanii* strains were either grown in rich medium (YPC-medium) for transformations or selective CA-medium (55) supplemented with an expanded trace element solution (CAB-medium) (31) for experiments. The strain with the hemagglutinin (HA)-tag on SepF, HTQ236, and H26 required addition of 0.45 mM uracil when grown in CAB-medium. The SepF depletion strain HTQ239 needed addition of 0.25 mM tryptophan for normal growth in CAB-medium and 0.45 mM uracil. To deplete SepF, strain HTQ239 was grown in CAB-medium in the absence of tryptophan. Cultures smaller than 5 ml were grown in 15 ml culture tubes rotating at 45 °C while larger cultures were grown in flasks on shakers set to 120 rpm at 45 °C. Plates for the growth of *H. volcanii* on solid medium after transformation were prepared as described previously by Allers et al. (55). Plates were incubated in plastic boxes to prevent evaporation at 45 °C. Growth curves of 20 ml cultures were automatically measured using the cell growth quantifier (CGQ) from aquila biolabs GmbH at 45 °C and constant shaking at 120 rpm. *Escherichia coli* strains were grown in LB-medium (56), supplemented with appropriate antibiotics (100 µg/ml ampicillin; 25 µg/ml kanamycin; 30 µg/ml chloramphenicol) if necessary, at 37 °C under constant shaking. Archaeal and bacterial strains used in this study are listed in supplementary table S1.

### Bioinformatic analysis of the putative archaeal SepF protein

The map of the genetic neighborhood of *hvo_0392* was drawn with Gene Graphics (57) with a region size of 6500 base pairs (bp). To obtain a general overview of the archaeal SepF proteins, sequences from selected bacterial SepF proteins were aligned with the putative SepF proteins from different archaea, using the MUSCLE (MUltiple Sequence Comparison by Log-Expectation (58, 59)) web service with default settings accessed via Jalview (Version 2.11.0) (60). The schematic overview of the putative SepF protein of *H. volcanii* was drawn with the Illustrator for Biological Sequences (IBS) (61). Amino acids with a conservation score > 7 in the alignment were indicated in the protein scheme. A comprehensive overview of the presence of SepF in prokaryotes was generated by a maximum-likelihood tree with the Molecular Evolutionary Genetics Analysis program (Mega, Version 10). The protein sequences were obtained by PSI-BLAST (Position-specific Iterated-Basic Local Alignment Search Tool) based on protein sequences from *B. subtilis* (WP_015483208.1), *C. glutamicum* (WP_040967618.1), *Synechocystis sp*. (WP_009632573.1), *H. volcanii* (ADE03140.1) and *Candidatus Woesearchaeota* (MBU90409.1). For each search six iterations were performed. Redundant sequences were removed with Jalview with a threshold of 75 % for archaeal sequences and 35 % for bacterial sequences. The obtained sequences were merged and aligned using MUSCLE with default settings from Mega-X. Badly aligned sequences were removed manually. The final alignment was then used to generate a maximum-likelihood tree with Mega-X and default settings.

### Plasmid construction

Enzymes for the amplification of inserts by polymerase chain reaction (PCR) (using PHUSION® polymerase), digestion and ligation were obtained from New England Biolabs (NEB). Plasmids for expression of fluorescently tagged proteins in *H. volcanii* were constructed via classical restriction enzyme-based cloning into plasmid pIDJL-40 or pTA1392 for single protein expression and pSVA3943 for double expression, following the manufacturer’s protocols. Inserts were amplified from genomic DNA of strain H26, isolated as described before (32). Genes were under control of a tryptophan inducible promotor *p*.*tnaA* (62).

For the generation of the plasmid to add an HA-tag to the genomically encoded *sepF (hvo_0392)* at its C-terminus two fragments were amplified: A fragment ∼500 bp up-stream of the *sepF* stop-codon with a 5’ KpnI restriction site and 3’ *ha* sequence with a stop-codon and a second fragment ∼500 bp down-stream of the *sepF* stop-codon with the complementary *ha* + stop-codon sequence at 5’ end and a XbaI restriction site at the 3’ end. Both fragments were assembled via overlap PCR using the forward primer of the up-stream fragment and the reverse primer of the down-stream fragment. The generated fragment was subsequently cloned into integrative plasmid pTA131.

The *sepF* knock-out plasmid was created by amplification of ∼500 bp of the up-stream region and ∼500 bp of the down-stream region of *sepF*. Both PCR products were linked together by digestion with BamHI and subsequent ligation. The fragment was then cloned into plasmid pTA131 via KpnI and XbaI. To generate the plasmid for the construction of the *sepF* depletion strain, the gene *sepF* was cloned in pTA1369, resulting in tryptophan-inducible *sepF*. The gene construct was subsequently cut out from pTA1369 with BglII and cloned in between the up- and down-stream regions of *sepF* in its knock-out plasmid opened with BamHI.

The plasmids for heterologous protein expression in *E. coli* were created with the FX-cloning system described by Geertsma and Dutzler (63). The sequences for the forward and the reverse primers to amplify *sepF, ftsZ1* (*hvo_0717*) and *ftsZ2* (*hvo_0581*) for FX-cloning were obtained from https://www.fxcloning.org/.

Primers used for the amplification of the different DNA fragments and the respective restriction enzymes for cloning are indicated in supplementary table S3. Plasmids used in this study are listed in supplementary table S2.

### Transformation of plasmids and genetic manipulation of *Haloferax volcanii*

*Haloferax volcanii* was transformed using polyethylene glycol 600 (PEG600) as described before (32). The selection for successful transformation was based on the complementation of the uracil auxotrophy of the used strains by the transformed plasmids. In brief: When 10 ml of a culture reached an optical density (OD_600_) of 0.8, cells were harvested (3000 g for 8 min) and spheroblasts were formed by resuspending in buffered spheroblasting solution (1 M NaCl, 27 mM KCl, 50 mM Tris-HCl (pH 8) and 15 % [w/v] sucrose) with 50 mM EDTA (pH 8). The resuspended cells were incubated at room temperature (RT) for 10 min. In the meantime, 1 µg of a respective demethylated plasmid, passed through a dam-/dcm-*E. coli* strain (NEB) before, was mixed with 83 mM EDTA and unbuffered spheroblasting-solution (1 M NaCl, 27 mM KCl and 15 % [w/v] sucrose) to a final volume of 30 µl. DNA was added to the spheroblasts and gently mixed. After 5 min of incubation an equal amount of 60 % PEG 600 was added, gently mixed and incubated for 30 min. Subsequently, 1.5 ml spheroblast-dilution solution (23 % salt water, 15 % [w/v] sucrose and 3.75 mM CaCl2) was added, the tubes were inverted and incubated for additional 2 min. After spheroblasts were harvested at 3000 g for 8 min, 1 ml regeneration solution (18 % salt water, 1 x YPC, 15 % [w/v] sucrose and 3 mM CaCl2) was added and the pellet transferred to a sterile 5 ml tube. The pellet was incubated standing for 1.5 h at 45 °C before it was resuspended by gentle shaking and additional incubation for 3.5 h rotating at 45 °C. Transformed cells were then harvested (3000 g for 8 min) and resuspended in 1 ml transformant-dilution solution (18 % salt water, 15 % [w/v] sucrose, 3 mM CaCl2). At last, 100 µl of 10 ^0^, 10 ^−1^ and 10 ^−2^ dilutions of the transformed cells were plated on selective plates.

### Integration of a genomic HA-tag at the *sepF locus*

For the integration of an endogenous HA-tag at *sepF* the pop-in / pop-out method was used (64). H26 was transformed with integrative plasmid pSVA3947 and subsequently grown on selective CA-plates until colonies were visible. To induce the pop-out, one transformant colony was transferred to 5 ml non-selective medium (YPC-medium) and grown at 45 °C. When the culture reached OD_600_ 1 it was diluted 1:500 into fresh YPC-medium. This was repeated three times in total. To select for successful pop-out events, 100 µl of the 10 ^−2^ diluted pop-out culture was plated on 50 µg/ml 5-fluoorotic acid (5-FOA) containing CA-plates. Colonies from the 5-FOA plate were screened via colony PCR for successful HA-tag integration. For the colony PCR material from single colonies was transferred into 300 µl dH_2_O and incubated for 10 min at 75 °C under constant shaking for complete cell lysis. Subsequently, 1 µl of the lysed cells were used as template and the PCR product sent for sequencing.

### Construction of a SepF depletion strain

To exchange the native promotor of *sepF* with a tryptophan-inducible promotor, integrative plasmid pSVA3954 was transformed in *H. volcanii* strain H98. Transformants were checked for correct up-stream integration and orientation of the construct by colony PCR. Subsequent pop-out procedure was performed as described before. To screen for the promotor exchange, pop-out colonies were first re-streaked on selective plates containing tryptophan and uracil. After colonies had formed, they were transferred to a selective plate with uracil but without tryptophan. Colonies that were not able to grow without tryptophan were checked via colony PCR and subsequent sequencing for successful promotor exchange.

### Microscopy

To investigate the cellular localization of the putative archaeal SepF protein alone or in combination with FtsZ1 or FtsZ2, fluorescence microscopy was used. To image *H. volcanii*, a 3 µl sample of early exponentially growing cells (OD_600_ < 0.1) was spotted on a 0.3 % [w/v] agarose pad containing 18 % buffered salt water (144 g/l NaCl, 18 g/l MgCl _2_ * 6 H_2_O, 21 g/l MgSO _4_ * 7 H_2_O, 4.2 g/l KCl, 12 mM Tris/HCl, pH 7.5). After the sample was dried on the pad it was covered with a cover slide and observed with an inverted microscope (Zeiss Axio Observer.Z1) equipped with a temperature-controlled chamber at 45° C.

For over-night time lapse microscopy, an agarose pad containing nutrients was prepared by mixing 0.3 % agarose [w/v] with CAB-medium supplemented with 5 % [w/v] sucrose. One ml of the nutrition pad solution was poured in a round Delta T Dish (Bioptechs Inc.). After the pad had solidified 3 µl of cells in very early exponential phase (OD_600_ 0.05-0.1) were dropped on the pad. After the spots were dried the whole pad was flipped up-side down into another Delta T Dish and the dish was closed with a lid (Bioptechs Inc.) to avoid evaporation. Cells were imaged at 45 °C over-night with image acquisition every 15 or 30 min for 16 h. For the observation of microcolonies, cells were grown over night on agarose pads containing nutrients at 45°C and were imaged the next morning.

To image strain HTQ239 transformed with different plasmids, precultures of the respective transformants were grown in 30 ml CAB-medium supplemented with tryptophan to an OD_600_ of 0.01. Cells were subsequently harvested at 3000 g for 20 min in a centrifuge heated to 45 °C. The supernatant was discarded, and the cells resuspended in pre-warmed CAB-medium and grown again at 45 °C under constant shaking. Cells were imaged every 3 h for 9 h and one last time 24 h after growing in media without tryptophan. Images were analyzed using FIJI (65) and the MicrobeJ plug-in (66).

### Heterologous expression and purification of SepF-His, FtsZ1-His and FtsZ2-Strep

The different proteins were produced by heterologous expression in the *E. coli* Rosetta™ strain. For overexpression, Rosetta™ cells were transformed with plasmid pSVA3969 (SepF-His), pSVA3970 (FtsZ1-His) or pSVA5912 (FtsZ2-Strep) and grown in 1 l LB-medium supplemented with kanamycin (25 µg/ml) and chloramphenicol (30 µg/ml) at 37 °C under constant shaking. After reaching an OD_600_ of 0.5 expression was induced by adding 0.5 mM β-D-1-thiogalactopyranoside (IPTG) (Fisher Science) and cells were grown for additional 3 h at 37 °C. Subsequently, cells were harvested by centrifugation at 6000 g for 20 min. The pellet was immediately frozen in liquid nitrogen and stored at −80 °C until used.

For purification of His-tagged proteins the frozen pellet was thawed and resuspended in 20 ml Buffer A (1.5 M KCl, 100 mM NaCl, 50 mM L4-(2-hydroxyethyl)-1-piperazineethanesulfonic acid (HEPES), 1 mM dithiothreitol (DTT), pH 7.5) supplemented with 10 µg/ml DNase I (Roche) and cOmplete™ EDTA-free Protease Inhibitor (Roche). Cells that produced SepF-His were opened using a Microfluidizer^®^, cells containing overproduced FtsZ1-His were lysed by sonication (Sonopuls, Bandelin; Probe KE76; 35 % power, 5 s pulse 5 s pause for 10 min). Cell debris were then removed by centrifugation at 3000 g for 15 min at 4 °C and the supernatant used for a second centrifugation step at 26000 g for 20 min at 4 °C. The soluble fraction of SepF-His or FtsZ1-His was manually loaded on a, with Buffer A pre-equilibrated, 5 ml HisTrap™ HP column (GE Healthcare). For further purification steps the column was connected to a liquid chromatography system (Azura, Knauer). The column was washed in three steps with 10 ml Buffer A containing 10 mM, 20 mM and 30 Mm imidazole, respectively. The protein was eluted from the column via an imidazole gradient from 30-400 mM imidazole in Buffer A and collected in 2 ml fractions.For the purification of FtsZ2-Strep, cells were lysed as described for FtsZ1-His. The soluble fraction was loaded on a 3 ml gravity column of Strep-Tactin® Sepharose® (iba-lifesciences). Subsequently the column was washed with 15 ml Buffer A. To elute the protein 3 ml of Buffer A supplemented with 2.5 mM desthiobiotin was added and the flow-through collected in 1 ml fractions. FtsZ1-His and FtsZ2-Strep were concentrated by ultrafiltration (Amicon® Ultra 4 ml, 3 kDA cut-off) and subsequently frozen in liquid nitrogen and stored at −80 °C.The purified SepF-His was used for size exclusion chromatography (see: Determination of the oligomeric state of the archaeal SepF) while purified FtsZ1 (130 µM) and FtsZ2 (62 µM) were used as positive controls for the immunoprecipitation experiments with HA-tagged SepF.

### Immunoprecipitation of HA-tagged SepF

*Haloferax volcanii* strain HTQ236 and H26 as control were grown in 333 ml CAB-medium supplemented with uracil to an OD_600_ of 0.5. The cells were harvested at 3000 g for 20 min and subsequently resuspended in 30 ml 18 % salt water buffered with 10 mM HEPES. For crosslinking 1.2 % [v/v] formaldehyde was added, and the cells incubated for 10 min at RT under constant shaking. To quench the crosslinking reaction 20 ml of 1.25 M glycine dissolved in 18 % salt water were added and the cells harvested as described before. Following, the cells were washed two times with 20 ml ice cold 18 % salt water with 1.25 M glycine. After the last washing step cells were resuspended in 3 ml 1 x phosphate-buffered saline (PBS) (137 mM NaCl, 2.7 mM KCl, 10 mM Na_2_HPO_4_, 1.8 mM KH_2_PO_4_, adjusted to pH 7.2) containing additionally 10 mM MgCl_2_ and 10 µg/ml DNase I (Roche) and lysed via sonication (Sonopuls, Bandelin; Probe MS73; 35 % power, 5 s pulse 5 s pause for 5 min). The lysed cells were then centrifuged at 3000 g and 4 °C for 15 min to remove cell debris. The supernatant was used for ultra-centrifugation to separate the membrane and cytosolic fraction: The samples were centrifuged at 200000 g and 4 °C for 2 h. Subsequently, the membrane fraction was resuspended in 1 x PBS supplemented with 1 % Triton-X [v/v] with a volume equal to the volume of the supernatant. Both fractions were used for immunoprecipitation. The HA-tagged proteins were captured with the Pierce™ HA-Tag Magnetic IP/Co-IP Kit (Thermo Scientific™). Magnetic beads (0.25 mg) were prepared according to the manufacturer’s protocol. The membrane and the cytosolic fractions respectively were added in 1 ml steps to the beads. Each loading step was followed by 30 min incubation at 4 °C while slowly rotating. The washing steps and the elution from the beads with non-reducing sample buffer were executed as described in the manufacturer’s protocol.

Pulldown fractions were either analyzed by mass spectrometry (MS) or Western blot analysis. For Western blotting the respective samples were separated by sodium dodecyl sulfate-polyacrylamide gel electrophoresis (SDS-PAGE). Gels were stained with Fairbanks (25 % [v/v] isopropanol, 10 % [v/v] acetic acid, 0.05 % [w/v] Coomassie R) staining solution or blotted on polyvinylidenfluorid (PVDF) membranes using the Trans-Blot Turbo Transfer System (BioRad). As positive controls purified FtsZ1 or FtsZ2 was loaded on the respective gel. Each antibody against FtsZ1 or FtsZ2 (received from Iain Duggin, Sydney) raised in rabbits was incubated with the membrane over-night in a 1:1000 dilution (in 1 x PBS + 0.1 % Tween-20) at 4 °C. Subsequently, the membrane was incubated with the secondary antibody against rabbit coupled with a horseradish peroxidase (HRP) (Sigma-Aldrich) for 1 h in a 1:10000 dilution. Signals from the Western blots were acquired with the iBright FL1500 imaging system (Invitrogen) after adding substrate for the HRP (Clarity Max Western ECL Substrate, BioRad) to the membrane. The MS sample preparation, measurement and data evaluation were performed by the Core Facility Proteomic of the ZBSA (Zentrum für Biosystemanalyse, Freiburg, Germany).

### Determination of the oligomeric state of the archaeal SepF

To determine the oligomeric state of SepF, 500 µl of freshly purified protein was loaded on a Superdex 200 10/300 GL column (GE Healthcare) equilibrated with Buffer A. Data was collected for 60 min at a constant flow-rate of 0.5 ml/min. Additionally, a gel filtration standard (#1511901, BioRad) was run under the same conditions as SepF. The eluted protein was concentrated by ultrafiltration (Amicon® Ultra 4 ml, 3 kDA cut-off) and subsequently analyzed by SDS-gel electrophoresis and Western blotting. The HRP-coupled anti-His antibodies (Abcam®) for the detection of the His-tag on SepF were incubated with the membrane overnight in a 1:10000 dilution.

## Acknowledgements

We acknowledge support from “Zentrales Innovationsprogramm Mittelstand (ZIM) des Bundesministeriums für Wirtschaft und Energie (BMWi) (grant number ZF4653901AJ8) and VW Momentum (Grant number 94933) for PN. The antibodies against FtsZ1 and FtsZ2 were a gift from Iain Duggin, UTS, Australia.

## Notes

### Competing Interest Statement

The authors have declared no competing interest.

