## Supplementary material for "Archaeal SepF is essential for cell division in *Haloferax volcanii*"

**Movie S1 Time lapse of depletion of SepF**

**Movie S2 Localization of SepF-GFP**

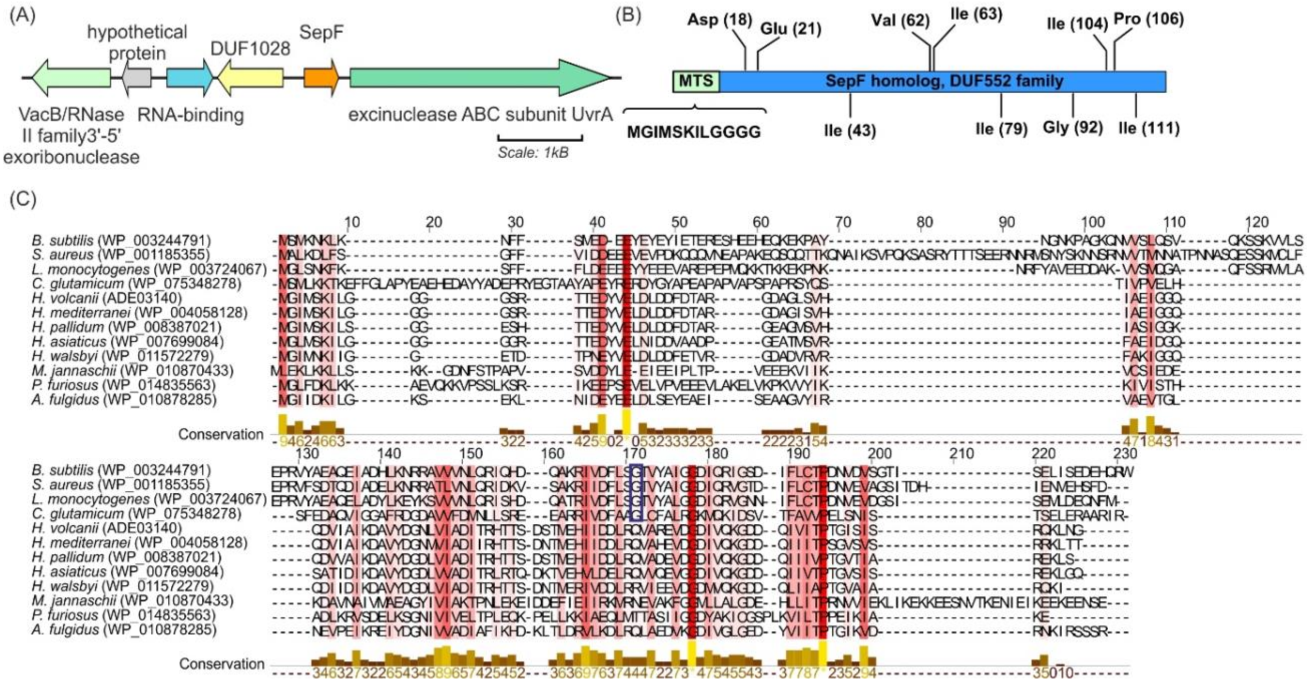


**Figure S1: Bioinformatical analysis of putative archaeal SepF proteins.** (A) Genetic-neighborhood map of *hvo_0392*. Mapped are 6500 bp of the *hvo_0392* region. The UvrA protein is a DNA-interacting ATPase and part of the UvrABC repair system that recognizes DNA lesions (1). The closest up-stream gene of *hvo_0392* encodes for a protein with a domain of unknown function (DUF1028). According to the Pfam database (2) some proteins of this family are associated with peptidoglycan binding. The following up-stream genes encode for an RNA-binding protein, a hypothetical protein and a VacB/RNase family II 3’-5’ exoribonuclease. (B) Schematic overview of the putative SepF protein from *H. volcanii*. The sequence of the membrane targeting sequence (MTS) (green) is indicated below and residues conserved between archaea and bacteria are highlighted. (C) Aligned protein sequences of bacterial SepFs with putative archaeal SepF proteins. Conserved residues are highlighted in a red color gradient from light red (conservation score of 6) to dark red (completely conserved). Residue G109 is indicated by a blue box. The conservation scores per site are indicated below the alignment.

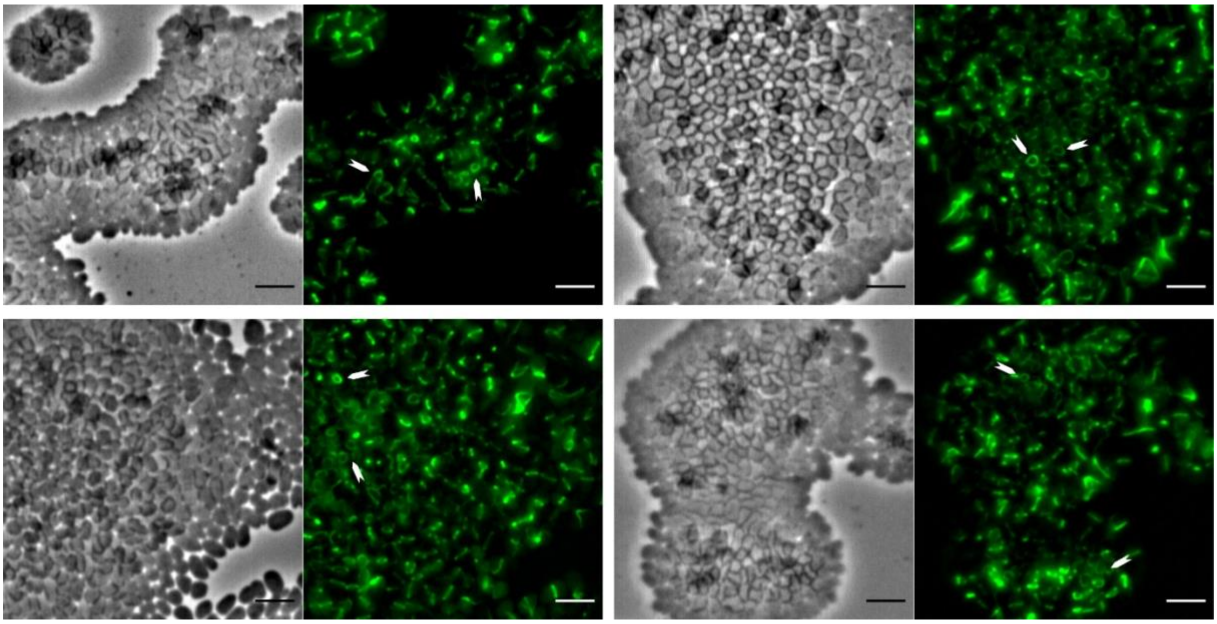

**Figure S2: SepF forms ring-like structures.** Microscopy pictures of H26 expressing SepF-GFP from plasmid pSVA2942 grown into microcolonies on agarose pads with nutrients. In tilted cells formation of SepF-GFP into a ring like structures is observable indicated by white arrows. Scale bar: 4 µm

(A)

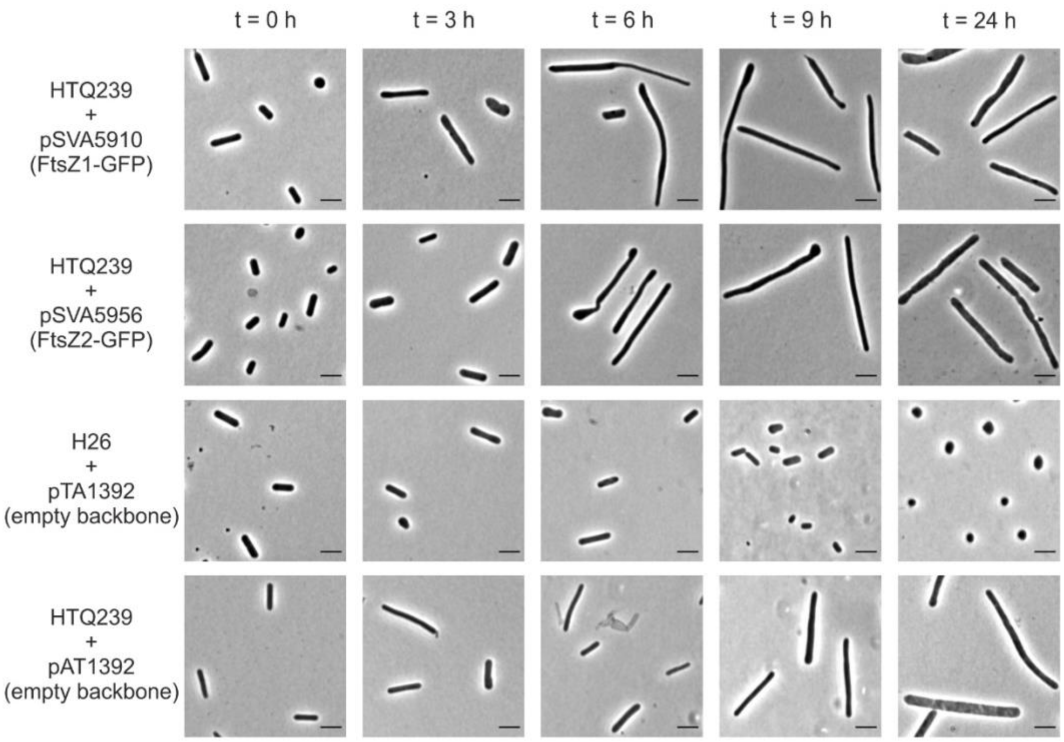


(B)

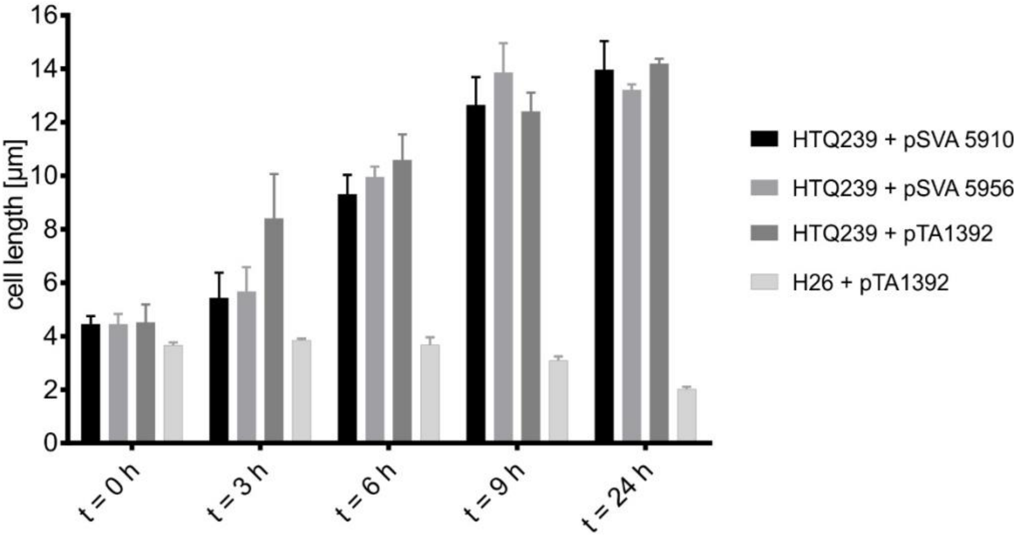


**Figure S3: Presence of a plasmid results into filamentation of HTQ239.** (A) Microscopy pictures of HTQ239 containing either FtsZ1-GFP or FtsZ2-GFP expression plasmids and pictures of H26 and HTQ239 containing empty backbone plasmid pTA1392 at different time points after SepF depletion. (B) Mean cell length of strainHTQ239 + pSVA5910, HTQ239 + pSVA5956 H26 + pTA1392 and HTQ239 + pTA1392 0, 3, 6, 9 and 24 h after SepF depletion. Cell length was obtained from three independent experiments including the cell size of > 700 cells per strain and time point.

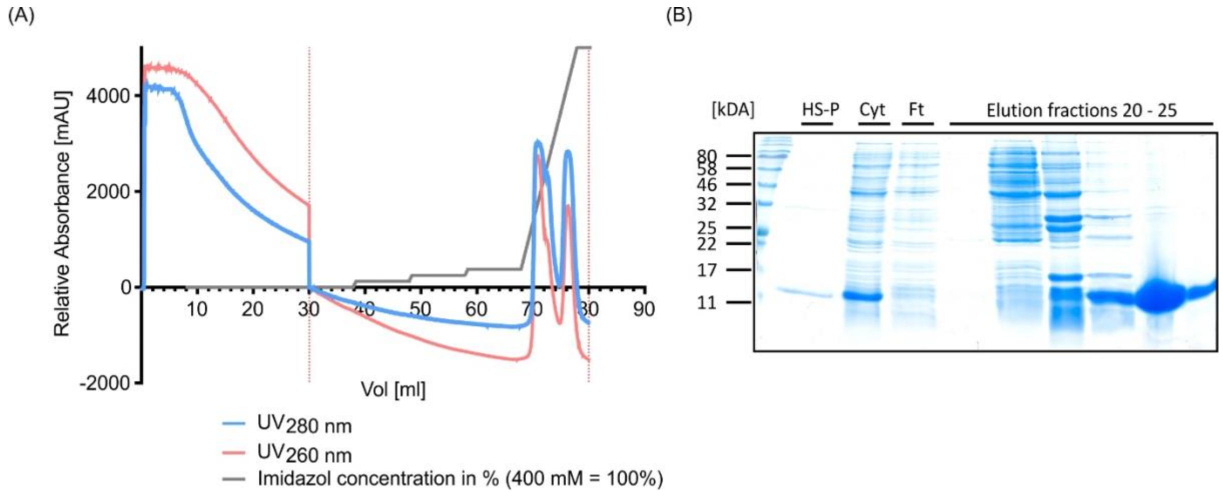


**Figure S4: SepF.** (A) Chromatogram of the purification measured at 280 nm (blue) and 260 nm (red). The imidazole gradient is indicated by a grey line. (B) SDS-gel of the purification showing the pellet fraction of the high spin centrifugation step (HS-P), the cytosolic fraction (Cyt) used for purification the flow-through (Ft) and the elution fractions 20-25 (10 µl samples per pocket).

**Table S1: Strains used in this study**

| Strain Name | Background strain | Genotype | Source/reference |
|---|---|---|---|
| ***H. volcanii*** | | | |
| H26 | - | Δ*pyrE2* | |
| H98 | - | Δ*pyrE2* Δ*hdrB* | (3) |
| HTQ236 | H26 | Δ*pyrE2 hvo_0392*::[*hvo_0392-ha*] | This study |
| HTQ239 | H98 | Δ*pyrE2* Δ*hdrB hvo_0392*::[*p.tnaA-hvo_0392-hdrB*$^{+}$] | This study |
| ***E. coli*** | | | |
| 10-beta Competent Cells "TOP10" | - | Δ*(ara-leu) 7697 araD139 fhuA* Δ*lacX74 galK16 galE15 e14-* ϕ*80*d*lacZ*Δ*M15 recA1 relA1 endA1 nupG rpsL* (Str$^{R}$) *rph spoT1* Δ*(mrr-hsdRMS-mcrBC)* | New England Biolabs |
| *dam*$^{-}$/*dcm*$^{-}$ Competent | - | *ara-14 leuB6 fhuA31 lacY1 tsx78 glnV44 galK2 galT22 mcrA dcm-6 hisG4 rfbD1 R(zgb210::Tn10)* Tet$^{S}$ *endA1 rspL136* (Str$^{R}$) *dam13::Tn9* (Cam$^{R}$) *xylA-5 mtl-1 thi-1 mcrB1 hsdR2* | New England Biolabs |
| Rosetta™(DE3) Competent Cells | - | F$^{-}$ *ompT hsdS*$_{B}$(r$_{B}^{-}$ m$_{B}^{-}$) *gal dcm* (DE3) pRARE (Cam$^{R}$) | Novagen |

**Table S2: Plasmids used in this study**

| Plasmids | Description | Primers used | Enzymes used | Source/reference |
|---|---|---|---|---|
| pTA131 | Integrative plasmid with a *pyrE2* selection marker for knock-outs in *H. volcanii* (Amp$^{r}$) | - | - | (3) |
| pIDJL-40 | Plasmid for the expression of proteins with a C-terminal GFP-tag and a *pyrE2, hdrB* selection markers (Amp$^{r}$) | - | - | (4) |
| pIDJL-114 | Plasmid for the expression of proteins with a C-terminal mCherry-tag and *pyrE2, hdrB* selection markers (Amp$^{r}$) | | | Iain Duggin |
| pTA1392 | Plasmid for the expression of proteins in *H. volcanii* under control of *p.tnaA* and *pyrE2, hdrB* selection markers (Amp$^{r}$) | | | (5) |
| pTA1369 | Integrative plasmid with a *pyrE2* and *hdrB* selection marker containing the *p.tnaA1* promotor to generate a | - | - | (6) |

| | | | | |
|---|---|---|---|---|
| | tryptophan inducible allele of a gene<br>($Amp^r$) | | | |
| pTA1992 | Plasmid for the constitutive expression of proteins in *H. volcanii* under control of as synthetic promotor (*pSyn)* with *pyrE2, hdrB* selection markers<br>($Amp^r$) | | | (6) |
| p7XC3H | FX cloning *E. coli* expression vector with T7 promoter and C-terminal 3C protease cleavage site and 10x His-tag<br>($Kan^r$) | - | - | (7) |
| p7XC3S | FX cloning *E. coli* expression vector with T7 promoter and C-terminal 3C protease cleavage site and Strep-tag<br>($Kan^r$) | | | (7) |
| pSVA3922 | Plasmid for the expression of proteins with a N-terminal GFP-tag and *pyrE2, hdrB* selection markers<br>($Amp^r$) | | | (8) |
| pSVA3933 | integrative plasmid with a *pyrE2* selection marker to generate a *hvo_0392* knock-out mutant<br>($Amp^r$) | 8025,<br>8026;<br>8027,<br>8028 | KpnI,<br>BamHI;<br>BamHI,<br>XbaI | This study |
| pSVA3943 | Plasmid to express two proteins, one with a c-terminal GFP tag and on with a c-terminal mCherry tag under the control of *p.tnaA.*<br>*pyrE2* selection marker based on pTA1392<br>($Amp^r$) | 7010<br>7011<br><br>8062,<br>8063 | NheI, EcoRI<br><br><br>BamHI,<br>NotI | This study |
| pSVA3942 | Tryptophan inducible SepF-GFP expression plasmid with a *pyrE2* selection marker<br>($Amp^r$) | 8066,<br>8067 | NdeI,<br>BamHI | This study |
| pSVA3947 | Integrative plasmid with a *pyrE2* selection marker to integrate an endogenous HA-tag at the C-terminus of SepF<br>($Amp^r$) | 8068.<br>8069;<br>8070,<br>8071 | KpnI, XbaI | This study |
| pSVA3951 | Tryptophan inducible GFP-SepF expression plasmid with a *pyrE2* selection marker<br>($Amp^r$) | 8083,<br>8084 | NheI,<br>BamHI | This study |
| pSVA3954 | Integrative plasmid for the exchange of *hvo_0392* native promoter<br>($Amp^r$) | 8096;<br>8097 | NdeI,<br>EcoRI;<br>BglII,<br>BamHI | This study |

| | | | | |
|---|---|---|---|---|
| pSVA3968 | Tryptophan inducible SepF-GFP, FtsZ1-mCherry double expression plasmid with a *pyrE2* selection marker ($Amp^r$) | 8092; 8093<br><br>8096, 8802 | EcoRI, BamHI (for FtsZ1)<br><br>NheI, NdeI (for SepF) | This study |
| pSVA3969 | Plasmid for expression of SepF with a C-terminal 10 x His-tag in *E. coli* based on p7XC3H ($Kan^r$) | 8803, 8804 | SapI | This study |
| pSVA3970 | Plasmid for expression of FtsZ1 with a C-terminal 10 x His-tag in *E. coli* based on p7XC3H ($Kan^r$) | 8805, 8806 | SapI | This study |
| pSVA5910 | Plasmid for expression of FtsZ1-GFP under the control of its native promotor ($Amp^r$) | 9703, 9704 | ApaI, BamHI | This study |
| pSVA5912 | Plasmid for expression of FtsZ2 with a C-terminal Strep-tag in *E. coli* based on p7XC3S ($Kan^r$) | 9705, 9706 | SapI | This study |
| pSVA5913 | Tryptophan inducible SepF-GFP, FtsZ2-mCherry double expression plasmid with a *pyrE2* selection marker. Based on pSVA3968 ($Amp^r$) | 9707, 9708 | EcoRI, BamHI | This study |
| pSVA5954 | Tryptophan inducible $SepF_{\Delta MTS}$-GFP expression plasmid with a *pyrE2* selection marker ($Amp^r$) | 9789, 8067 | NdeI, BamHI | This study |
| pSVA5955 | Tryptophan inducible $MTS_{SepF}$-GFP expression plasmid with a *pyrE2* selection marker ($Amp^r$) | 11013, 11014 | NdeI, NotI | This study |
| pSVA5956 | Plasmid for expression of FtsZ2-GFP under the control of its native promotor with a *pyrE2* selection marker ($Amp^r$) | 9793, 9794 | ApaI/BamHI | This study |
| pSVA5960 | Plasmid for constitutive expression of SepF under the control of a synthetic promotor with a *pyrE2* selection marker ($Amp^r$) | 8867, 8024 | NcoI, NheI (insert); PciI, NheI (plasmid) | This study |

**Table S3: Primers used in this study**

| Primer Number/Name | Sequence 5'→3' | Description |
|---|---|---|

| | | |
|---|---|---|
| 7010 | CGGCTAGCATGAGTAAAGGAGAAGAACTTTTCACTGGAGTTGTCCCAATTCTTGTTG | Forward primer for the amplification of *gfp* with a NheI restriction site (cloned in pTA1392) |
| 7011 | GGAATTCTCACTTCTCGAACTGCGGGTGCGACCATTTGTATAGTTCATCCATGCCATG | Reverse primer for the amplification of *gfp* with an EcoRI restriction site (cloned in pTA1392) |
| 8024 | TCTAGCTAGCGCCGTTGAGCTTCTGC | Reverse primer for the amplification of *hvo_0392* with NheI restriction site (cloned in pIDJL-40 and pTA1992) |
| 8025 | GATAGGTACCAGCGAACGTCGCCTCGGAAC | Forward primer for the amplification of the up-stream region of *hvo_0392* with KpnI restriction site (cloned in pTA131) |
| 8026 | GTAGGATCCAGGACGTAGAGAGGCCACAAC | Reverse primer for the amplification of the up-stream region of *hvo_0392* with BamHI restriction site (cloned in pTA131) |
| 8027 | GTAGGATCCCGACGGCGGACGACACCC | Forward primer for the amplification of the down-stream region of *hvo_0392* with BamHI restriction site (cloned in pTA131) |
| 8028 | GTACTCTAGAATGTCCTGTGCGGACTGCTC | Reverse primer for the amplification of the down-stream region of *hvo_0392* with XbaI restriction site (cloned in pTA131) |
| 8062 | CGGGATCCCCAATGCATGTGAGCAAGGGCGAGGAGGATAAC | Forward primer for the amplification of *mcherry* with a BamHI and NsiI restriction site (cloned in pTA1392) |
| 8063 | ATAAGAATGCGGCCGCTTACTTGTACAGCTCGTCCATGCC | Reverse primer for the amplification of *mcherry* with a NotI restriction site (cloned in pTA1392) |
| 8066 | GGAATTCCATATGGGTATCATGAGTAAGATTCTCG | Forward primer for the amplification of *hvo_0392* with NdeI restriction site (cloned in pIDJL-40) |
| 8067 | CTAGGATCCGCCGTTGAGCTTCTGCC | Reverse primer for the amplification of *hvo_0392* with BamHI restriction site (cloned in pIDJL-40 and pSVA5954) |
| 8068 | GGGGTACCGACGCCCGTTGGAACCGCCGACT | Forward primer for the amplification of ~500 Bp up-stream region of the *sepF* stop codon with a KpnI restriction site |
| 8069 | TCACGCGTAGTCCGGGACGTCGTACGGGTAGCTGCCGCCGTTGAGCTTCTGCC | Reverse primer for the amplification of ~500 Bp up-stream region of the *sepF* stop codon containing the coding sequence for a HA-tag + stoop codon |
| 8070 | GGCAGCTACCCGTACGACGTCCCGGACTACGCGTGACGACGGCGGACGACACC | Forward primer for the amplification of ~500 Bp down-stream region of the *sepF* stop codon containing the coding sequence for a HA-tag + a stop codon (the HA tag sequence is complementary to the on in primer 8069) |

| | | |
|---|---|---|
| 8071 | TGCTCTAGAATGTCCTGTGC GGACTGCTC | Reverse primer for the amplification of ~500 Bp down-stream region of the *sepF* stop codon with a XbaI restriction site |
| 8055 | GCGCGTCGATGAGTCGTTTC | Forward primer to screen for successful HA-tag incorporation |
| 8083 | TCTAGCTAGCATCATGAGTAA GATTCTCGGTGGTG | Forward primer for the amplification of *hvo_0392* with NheI restriction site (cloned in pSVA3922) |
| 8084 | CTAGGATCCTCAGCCGTTGA GCTTCTG | Reverse primer for the amplification of *hvo_0392* with BamHI restriction site (cloned in pSVA3922) |
| 8087 | GTAGTCCGGGACGTCGTACG G | Reverse primer binding the HA-DNA sequence to check for successful tag incorporation in HTQ236 |
| 8092 | CCGGAATTCATGGACTCTATC GTCGGCGAC | Forward primer for the amplification of *ftsZ1* with EcoRI restriction site (cloned in pSVA3943) |
| 8093 | CGCGGATCCCTCGACGTAGT CGATGTCTTC | Reverse primer for the amplification of *ftsZ1* with BamHI restriction site (cloned in pSVA3943) |
| 8096 | GGAATTCCATATGGGTATCAT GAGTAAGATTCTCGG | Forward primer for the amplification of *hvo_0392* with NdeI restriction site (cloned in pTA1369 and pSVA3943) |
| 8097 | CCGGAATTCTCAGTGGTGGT GGTGGTGGTGGCCGTTGAGC TTCTGCCGC | Revers primer for the amplification of *hvo_0392* with a NdeI restriction site and 6 x His coding sequence (cloned in pTA1369) |
| 8802 | TCTAGCTAGCGCCGTTGAGC TTCTGC | Reverse primer for the amplification of *hvo_0392* with a NheI restriction site (cloned in pSVA3943) |
| 8803 | ATATATGCTCTTCTAGTGGTA TCATGAGTAAGATTCTCGGTG GT | Forward primer for the amplification of *hvo_0392* with SapI restriction site (cloned in p7XC3H) |
| 8804 | CCAAGTGCTCTTCATGCGCC GTTGAGCTTCTGCCGCGAGA TAGA | Reverse primer for the amplification of *hvo_0392* with SapI restriction site (cloned in p7XC3H) |
| 8805 | ATATATGCTCTTCTAGTGACT CTATCGTCGGCGACGCAATT GAC | Forward primer for the amplification of *ftsZ1* with SapI restriction site (cloned in p7XC3H) |
| 8806 | TATATAGCTCTTCATGCTTCG ACGTAGTCAATGTCTTCGAGA CG | Reverse primer for the amplification of *ftsZ1* with SapI restriction site (cloned in p7XC3H) |
| 8817 | TCGTGGCGAGAACGGAACAG | Forward primer binding *p.tnaA* for colony PCR on HTQ239 |
| 8818 | GGGCAGACGAACACGATGAC | Reverse primer binding *hvo_0392* for colony PCR on HTQ239 |
| 8867 | CATGCCATGGGTATCATGAG TAAGATTCTCGG | Forward primer for the amplification of *hvo_0392* with NcoI restriction site (cloned in pTA1992) |

| | | |
|---|---|---|
| 9703 | TTCGGGCCCAAGCTCTCGGC CGGGCAGTCC | Forward primer for the amplification of *ftsZ1* with its promotor region and an ApaI restriction site (cloned in pIDJL-40) |
| 9704 | CGCGGATCCCTCGACGTAGT CGATGTCTTC | Reverse primer for the amplification of *ftsZ1* with BamHI restriction site (cloned in pIDJL-40) |
| 9705 | ATATATGCTCTTCTAGTCAGG ATATTGTTCGCGAGGCGATG GAA | Forward primer for the amplification of *ftsZ2* with SapI restriction site (cloned in p7XC3S) |
| 9706 | TATATAGCTCTTCATGCCCGA ATGACGTCGAGACCGTTGTT CTT | Reverse primer for the amplification of *ftsZ2* with SapI restriction site (cloned in p7XC3S) |
| 9707 | CCGGAATTCATGCAGGATAT CGTTCGCGAG | Forward primer for the amplification of *ftsZ2* with EcoRI restriction site (cloned in pSVA3943) |
| 9708 | CGCGGATCCCCGGATGACGT CGAGACC | Reverse primer for the amplification of *ftsZ2* with BamHI restriction site (cloned in pSVA3943) |
| 9789 | GGAATTCCATATGGGTGGTG GTTCCCGAAC | Forward primer for the amplification of truncated *hvo_0392* ($SepF_{\Delta MTS}$) with NdeI restriction site (cloned in pIDJL-40) |
| 9793 | TTCGGGCCCTTTCGGGTGTA AGACACTCG | Forward primer for the amplification of *ftsZ2* with its promotor region and an ApaI restriction site (cloned in pIDJL-40) |
| 9794 | CGCGGATCCCCGGATGACGT CGAGACC | Reverse primer for the amplification of *ftsZ2* with BamHI restriction site (cloned in pIDJL-40) |
| 11013 | ATTGCGCATATG*GGTATCATG AGTAAGATTCTCGGTGGTGG TGGT*GGATCCATGAGTAAAG GAGAAGAAC | Forward primer for the amplification of gfp containing the gene *sequence for the MTS of SepF* and NdeI restriction site (cloned in pTA1392) |
| 11014 | ATAAGAATGCGGCCGCTTATT TGTATAGTTCATCCATGCCAT G | Reverse primer for the amplification of *gfp* with a NotI restriction site (cloned in pTA1392) |

**Table S4: MS results from pulldown experiments. Identified proteins in the pulldown fractions of whole cell lysate from H26 and HTQ236 are shown. FtsZ2 exclusively found in the pulldown fraction of HTQ236 is highlighted (yellow).**

| Protein IDs (Uniprot) | Protein names | Peptide counts (unique) | Sequence coverage [%] | PEP | Intensity Sample from 1 (H26) | Intensity Sample from 2 (HTQ236) |
|---|---|---|---|---|---|---|
| D4GP60 | Cobalt-factor-III C17-methyltransferase | 1 | 5.4 | 2.13E-22 | 413500 | 235420 |
| D4GP61 | Cobalamin biosynthesis protein CbiG | 1 | 4 | 0.0079128 | 44590 | 0 |
| D4GP63 | Cobalt-factor-II C20-methyltransferase | 1 | 7.7 | 7.08E-05 | 144240 | 255540 |
| D4GP64 | Probable cobalt-precorrin-6B C(15)-methyltransferase (decarboxylating) | 5 | 39.1 | 3.25E-59 | 951400 | 728940 |
| D4GQU5 | Class II aldolase (Homolog to L-fuculose-phosphate aldolase) | 1 | 4.3 | 1.20E-10 | 134800 | 0 |
| D4GRF0 | Cobyrinate a,c-diamide synthase | 3 | 10.9 | 5.62E-20 | 261230 | 628040 |
| D4GSH7 | Cell division protein FtsZ2 | 13 | 55.8 | 0 | 0 | 22549000 |
| D4GSW9 | Uridylate kinase | 1 | 7.6 | 3.29E-39 | 141920 | 146280 |
| D4GT59 | Succinate--CoA ligase [ADP-forming] subunit alpha | 2 | 9.3 | 3.63E-12 | 2417300 | 0 |
| D4GTI4 | Aconitate hydratase | 7 | 18.7 | 1.66E-122 | 381180 | 1842800 |
| D4GTJ3 | PRC domain protein | 2 | 37 | 5.28E-26 | 0 | 866450 |
| D4GTX8 | 50S ribosomal protein L19e | 5 | 33.1 | 3.71E-30 | 1559400 | 1037100 |
| D4GTY4 | 30S ribosomal protein S4e | 3 | 17 | 7.60E-45 | 198960 | 0 |
| D4GTY7 | 30S ribosomal protein S17 | 2 | 23.7 | 2.37E-86 | 869560 | 392600 |
| D4GU54;D4GZ05 | GalE family epimerase/dehydratase | 4;1 | 19.6 | 1.87E-243 | 620070 | 786560 |
| D4GU72 | Low-salt glycan biosynthesis protein Agl12 | 4 | 26.5 | 1.70E-76 | 588470 | 0 |
| D4GUM1 | Beta-lactamase domain protein | 2 | 4.2 | 2.03E-19 | 87886 | 0 |
| D4GW49 | Ribonuclease J | 3 | 9.6 | 1.80E-68 | 412660 | 443750 |
| D4GWR9 | Putative glycosyltransferase, type 1 | 1 | 4 | 0.00056575 | 171630 | 0 |
| D4GXC7 | UPF0173 metal-dependent hydrolase HVO_1290 | 1 | 6.3 | 7.88E-17 | 454890 | 262670 |
| D4GYD4 | D-gluconate dehydratase | 1 | 6.8 | 8.28E-06 | 0 | 143940 |
| D4GYN6 | Ornithine carbamoyltransferase | 2 | 11 | 1.07E-93 | 90063 | 141780 |
| D4GYP0 | Putative [LysW]-L-2-aminoadipate/[LysW]- | 5 | 26.2 | 1.20E-131 | 1070500 | 2207500 |

| | | | | | | |
|---|---|---|---|---|---|---|
| | L-glutamate phosphate reductase | | | | | |
| D4GYQ7 | ABC-type transport system periplasmic substrate-binding protein (Probable substrate dipeptide/oligopeptide) | 2 | 5.9 | 1.16E-15 | 260690 | 143520 |
| D4GZY6 | Elongation factor 1-alpha | 5 | 22.6 | 1.03E-148 | 517430 | 2306600 |
| D4GZY7 | 30S ribosomal protein S10 | 2 | 22.5 | 3.51E-08 | 0 | 181910 |
| D4H018 | Putative SepF protein | 6 | 80.5 | 7.29E-274 | 246670 | 74749000 |
| D4H036 | PRC domain protein | 3 | 42.3 | 2.74E-27 | 0 | 6214600 |
| O30560 | Thermosome subunit 2 | 3 | 11.5 | 8.73E-30 | 370730 | 0 |
| O30561 | Thermosome subunit 1 | 3 | 9.8 | 1.68E-126 | 291550 | 0 |
| Q48328 | DNA repair and recombination protein RadA | 4 | 15.7 | 5.80E-21 | 0 | 271570 |
| Q48333 | V-type ATP synthase beta chain | 12 | 38.2 | 2.74E-130 | 2810300 | 772780 |

References Supplementary Material


1. Grossman,L. and Yeung,A.T. (1990) The UvrABC endonuclease system of Escherichia coli--a view from Baltimore. *Mutat. Res.*, **236**, 213–221.
2. El-Gebali,S., Mistry,J., Bateman,A., Eddy,S.R., Luciani,A., Potter,S.C., Qureshi,M., Richardson,L.J., Salazar,G.A., Smart,A., *et al.* (2019) The Pfam protein families database in 2019. *Nucleic Acids Res.*, **47**, D427–D432.
3. Allers,T., Ngo,H.-P., Mevarech,M. and Lloyd,R.G. (2004) Development of additional selectable markers for the halophilic archaeon *Haloferax volcanii* based on the *leuB* and *trpA* genes. *Appl. Environ. Microbiol.*, **70**, 943–53.
4. Duggin,I.G., Aylett,C.H.S., Walsh,J.C., Michie,K.A., Wang,Q., Turnbull,L., Dawson,E.M., Harry,E.J., Whitchurch,C.B., Amos,L.A., *et al.* (2015) CetZ tubulin-like proteins control archaeal cell shape. *Nature*, **519**, 362–365.
5. Gamble-Milner,R. (2016) Genetic analysis of the Hel308 helicase in the archaeon Haloferax volcanii. *PhD thesis, Univ. Nottingham*.
6. Braun,F., Thomalla,L., van der Does,C., Quax,T.E.F., Allers,T., Kaever,V. and Albers,S.-V. (2019) Cyclic nucleotides in archaea: Cyclic di-AMP in the archaeon Haloferax volcanii and its putative role. *Microbiologyopen*, **8**, 1–23.
7. Geertsma,E.R. and Dutzler,R. (2011) A versatile and efficient high-throughput cloning tool for structural biology. *Biochemistry*, **50**, 3272–3278.
8. Nussbaum,P., Ithurbide,S., Walsh,J.C., Patro,M., Delpech,F., Rodriguez-Franco,M., Curmi,P.M.G., Duggin,I.G., Quax,T.E.F. and Albers,S.-V. (2020) An oscillating MinD protein determines the cellular positioning of the motility machinery in archaea. *bioRxiv*,

10.1101/2020.04.03.021790.
